# Schema cell formation in orbitofrontal cortex is suppressed by hippocampal output

**DOI:** 10.1101/2023.05.03.539307

**Authors:** Wenhui Zong, Jingfeng Zhou, Matthew P.H. Gardner, Zhewei Zhang, Kauê Machado Costa, Geoffrey Schoenbaum

**Affiliations:** Intramural Research Program of the National Institute on Drug Abuse, Baltimore, MD, USA; Chinese Institute for Brain Research, Beijing (CIBR), China; Concordia University, Montreal, Canada

**Keywords:** orbitofrontal cortex, hippocampus, schema, learning, single-unit, rat

## Abstract

Both orbitofrontal cortex (OFC) and hippocampus (HC) are implicated in the formation of cognitive maps and their generalization into schemas. However how these areas interact in supporting this function remains an open question, with some proposals supporting a serial model in which OFC draws upon task representations created by HC to extract key behavioral features and others proposing a parallel model in which both regions construct representations that highlight different types of information. Here we tested between these two models by asking how schema correlates in OFC would be affected by inactivation of HC output, after learning and during transfer across problems. We found the prevalence and content of schema correlates were unaffected by inactivation after learning, while inactivation during learning accelerated their formation. These results contradict a serial model and favor the proposal that OFC and HC operate in parallel to extract different features defining cognitive maps and schemas.

## Introduction

The orbitofrontal cortex (OFC) and hippocampus (HC) are both associated with the process of forming mental constructs – cognitive maps ^1,2^ – that permit adaptive behavior in situations where novelty or incomplete information prevents reliance on past experience ^3-5^. While first applied to explain the role of HC in mapping space and other informational dimensions in relational memory, the same term accurately describes the involvement of OFC in sussing out the components and relationships that define the world around us, particularly as relevant to our behavioral goals or purpose in a particular setting. Accurate knowledge of such task spaces – composed of the internally-specified states and state transitions that comprise the task at hand ^6,7^ – is enormously useful, whether navigating a maze to obtain pellets, a metro system to reach the airport, or social structures to get ahead. Having an accurate task map allows us to rapidly recognize new or incomplete information and then respond in a manner consistent with our needs and desires. This principle extends to the formation of schemas, which arguably represent generalized cognitive maps, and facilitate the transfer of knowledge to new problems of a type, as when knowledge of one metro system makes it easier to learn to use another. Neural activity reflecting schema formation has been demonstrated in both OFC and HC ^8-11^.

This convergence in function puts renewed emphasis on understanding how the two areas interact. Historically addressing this has been hampered by the very different tasks used to study HC, which typically focus on spatial information and navigation, versus those applied to OFC, which normally use non-spatial sensory modalities, especially chemosensory, in simpler Pavlovian or instrumental tasks. Notable exceptions to this dichotomy have shown that OFC maps spatial relationships in settings normally used to assess HC function ^12-15^ and that HC reflects information and contributes to adaptive behaviors more normally associated with OFC ^16-22^. When neural activity in the two areas is directly compared in the same task similarities and differences are evident ^10,11,23-28^; both areas construct a model of the task space, however the OFC appears to give precedence to biologically significant information, whereas HC represents externally-defined states with greater fidelity, even when incidental to task performance ^10,27,28^.

Yet while such comparative studies provide glimpses into how the two areas may interact in the formation of cognitive maps and schemas, they are usually conducted at steady state rather than during learning, rarely address transfer to new problems, and generally do not test for effects of manipulations of one area on correlates in the other. As a result, current evidence can be used to support either serial or parallel processing models ^10,27-33^. For instance, the HC may identify the true or externally valid task map or schema and then send it to areas such as OFC for extraction of behaviorally relevant features, both within and across problems. In this scenario, task representations in the OFC would be heavily dependent on HC processing. Alternatively, the OFC and HC may function in parallel, extracting different information relevant to task mapping and schema formation, according to each area’s unique functions. Under this arrangement, many features of the representations in the OFC would be independent of HC.

Here we tested between the predictions of these two models, asking specifically how generalized representations – schemas – encoded in single unit activity in the OFC are affected by inactivation of HC output, both during performance on established problems and during transfer to new problems. Our results distinguish between two alternative models, strongly favoring the proposal that OFC and HC operate in parallel to extract different features defining cognitive maps and schemas during the integration of new information.

## Results

Single-unit activity was recorded in OFC in rats (n’s = 4 female, 4 male) performing an odor sequence task built with a nested standard go/no-go odor discrimination (Figure 1A). In each trial, the rats sampled an odor presented at a centrally located port and then had to decide whether to respond at a nearby fluid well for a sucrose reward. Rather than being randomized, however, the odor cues presented on successive trials were arranged in a predictable, fixed sequence to define trajectories through a virtual “figure 8” maze. In initial training, ten different odor cues were arranged to form two unique figure 8 mazes, with similar reward structures (Figure 1B), and rats did two 80-trial blocks of each maze in each session. Critically, the rats could use the odor cues on each trial to correctly respond for reward, however they could also use the sequence to anticipate reward many trials into the future, like a rat traveling through a sequence of positions on an actual figure 8 maze.

**Figure 1.**
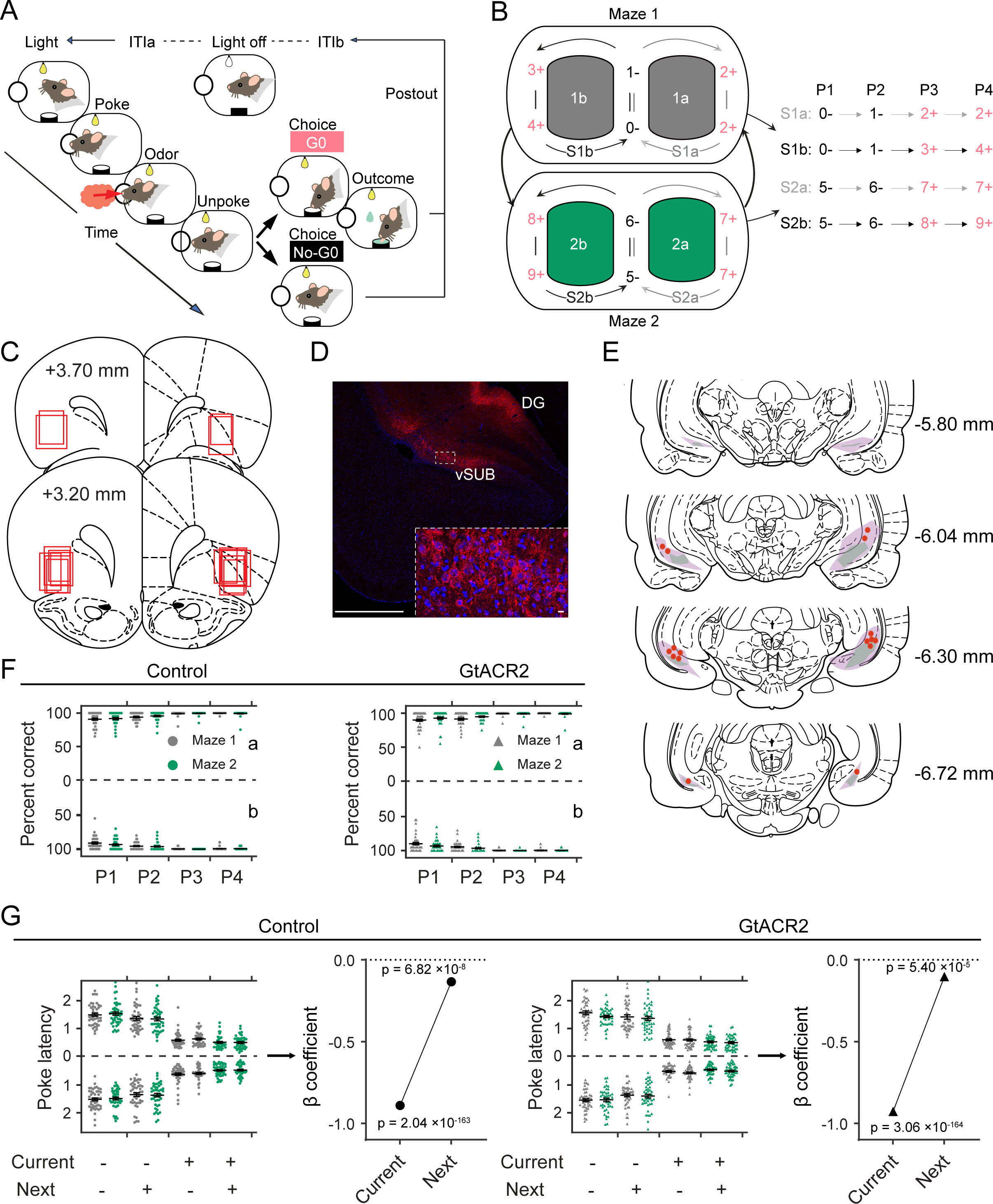
Task Design, Histology, and Behavior. (A) Schematic illustrating the events of a trial in the odor-sequence task. The illumination of two overhead houselights indicated the start of each trial. After poking into the central odor port and sampling the presented odor, rats could respond with a ‘‘go’’ to obtain a sucrose reward or a ‘‘no-go’’ to avoid a prolonged inter-trial interval. (B) Odor-sequence task is illustrated as two virtual figure 8 mazes. Ten odors were organized into two sequence pairs (S1 and S2), each comprising two subsequences (A and B). Each subsequence consists of four trials or positions (P1–P4) indicated by odor numbers. Red +, rewarded; black −, non-rewarded; 0–9, odor identities; arrows indicate sequence transitions. (C) Reconstruction of recording locations in lateral OFC. Approximate extent of recording locations in each rat is represented by red squares. (D) Virus expression. An AAV virus carrying the soma-targeted GtACR2-FusionRed construct under the CaMKIIa promoter was injected into the ventral subiculum bilaterally. GtACR2-expressing neurons were identified using immunohistochemistry. Red, GtACR2; Blue, DAPI. GtACR2-expressing neurons were found in the ventral subiculum (vSUB) and dentate gyrus (DG). Individual neurons expressing GtACR2 are magnified in the dashed white box. Scale bars indicate 1mm and 10µm, individually. (E) Reconstruction of GtACR2 expression and optical fiber placement. The maximal and minimal extent of GtACR2 expression is indicated by purple and green colors. Red dots indicate optical fiber placement. (F - G) Percent correct (F) and latency to poke into the odor port to initiate a trial after light onset (G) on each trial type in S1a, S2a (above *y*-axis), S1b, S2b (below *y*-axis) for control (left) and GtACR2 (right) sessions. Gray, Maze1; Green, Maze2. Error bars represent standard errors of the mean (SEM). Four-way ANOVAs confirmed significant main effects of position on both measures (% correct: F_1,408_ = 145.5, p = 4.2 × 10^-82^, η ^2^ = 0.24; poke latency: F = 889.1, p = 1.0 × 10^-323^, η ^2^ = 0.66), with reward driving more accurate and faster performance. Further regression analyses on the latency to initiate trials showed that this measure was affected by reward on both the current and next trials. Notably, in these analyses, there were no effects of inactivation (F’s < 0.82; p’s > 0.36; η ^2^’s < 0.0006). Error bars are SEM.

Once rats were trained to perform the task, electrodes were implanted in OFC to allow single-unit recording and fibers were implanted over ventral subiculum following infusion of pAAV-CKIIa-stGtACR2-FusionRed ^34^ (Addgene viral prep #105669-AAV1) to allow inactivation of hippocampal outflow (Figure 1C-E). Recording began 5-6 weeks after recovery from surgery and retraining on the task while tethered. During recording, each rat completed sessions in which 465nm light was delivered to activate GtACR2, and *inactivate* ventral subiculum, during each trial. Each inactivation session was followed by a reminder session and then a second recording session at the same location, during which light of an ineffective wavelength - 630 nm - was delivered to serve as a control ^35^. In both inactivation and control sessions, rats maintained highly accurate discrimination performance at all positions in both mazes (Figure 1F) and showed differences in their latencies to initiate trials, indicating use of the sequences to predict reward at the start of both the current and next trial (Figure 1G). There were no effects of maze or inactivation (see figure captions for statistics).

We recorded a total of 1856 units in OFC during the control sessions and 1834 units during the inactivation sessions. To visualize the patterns of firing during task performance, we calculated the activity of each single unit during each of 9 epochs tied to the specific events spanning each trial [ITIa, light, poke, odor, unpoke, choice, outcome, postOutcome, ITIb] for each of the 8 positions in each maze. This analysis revealed a great variety of patterns; however, the activity of individual units was generally influenced by some combination of trial epoch, reward, and position (Figures 2A-C). Overall, single unit activity in OFC was significantly influenced by each of these variables, with no apparent effect of inactivation (Figures 2D-F).

**Figure 2.**
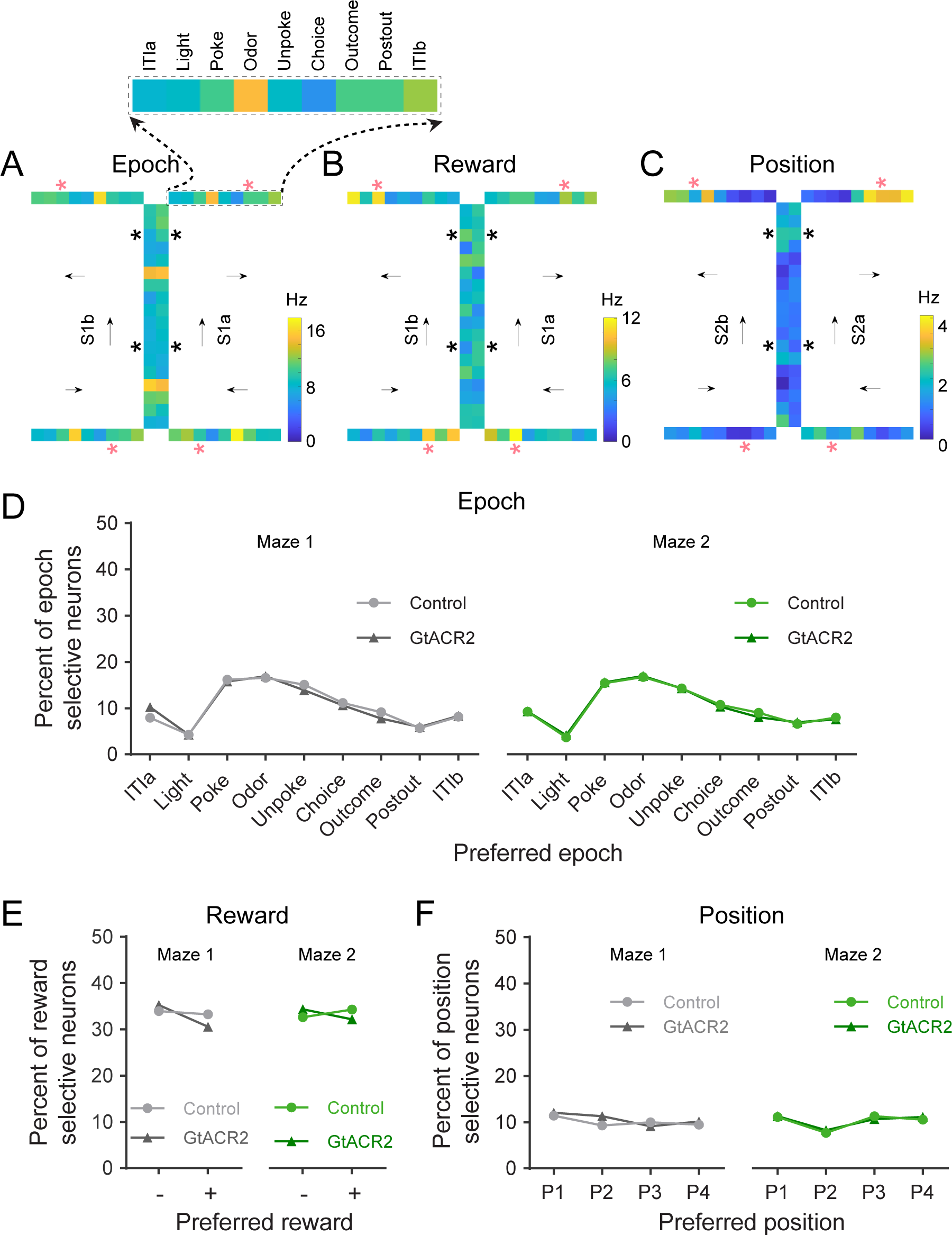
Exemplar units illustrating the influence of epoch, reward, position, and quantification across the population. (A - C) Heatmaps of OFC neurons showing epoch- (A), reward- (B), and position-specific (C) firing in the figure 8 task. In each panel, the heatmap shows average activity in each epoch at each position in one maze. Individual squares corresponding to each epoch are magnified in the black dashed box. Arrows represent sequence directions. Red * marks the reward epoch on rewarded trial types (P3 and P4), while black * marks the reward epoch for non-rewarded trial types (P1 and P2). (D - F) Plots show the percentage of the OFC neurons whose firing was significantly modulated by each factor (ANOVAs, p < 0.01), with each neuron assigned to the condition of maximal firing; there were no effects of inactivation (χ^2^’s < 1.42; p’s > 0.23; Chi-squared test). Error bars are SEM.

Importantly while the activity of some units discriminated between the two mazes (Figure S1), many showed very similar discrete firing patterns across them, consistent with representation of a generalized cognitive map or schema of the virtual figure 8 task. The generalization of the representations across the two mazes typically reflected the influence of the same variables noted above to impact unit firing, specifically trial epoch (Figure 3A), reward (Figure 3B), position (Figure 3C), or some combination of these factors (Figure 3D and see also Figures S2-3). While the generalization of variables such as epoch or reward would not necessarily require recognition of the common structure between the two mazes (e.g. examples in Figures. 3A and B), in other cases, generalization required recognition of this arbitrary structure. For example, in some units, activity was driven by the meaning of specific positions within the sequence (e.g. cell in Figure 3C, which fired most at P2 during choice->ITIb epochs), and in others it appeared to reflect still more idiosyncratic information characterizing particular epochs and positions (e.g. cell in Figure 3D, which fired at rewarded positions and at unpoke at P1 and P2).

**Figure 3.**
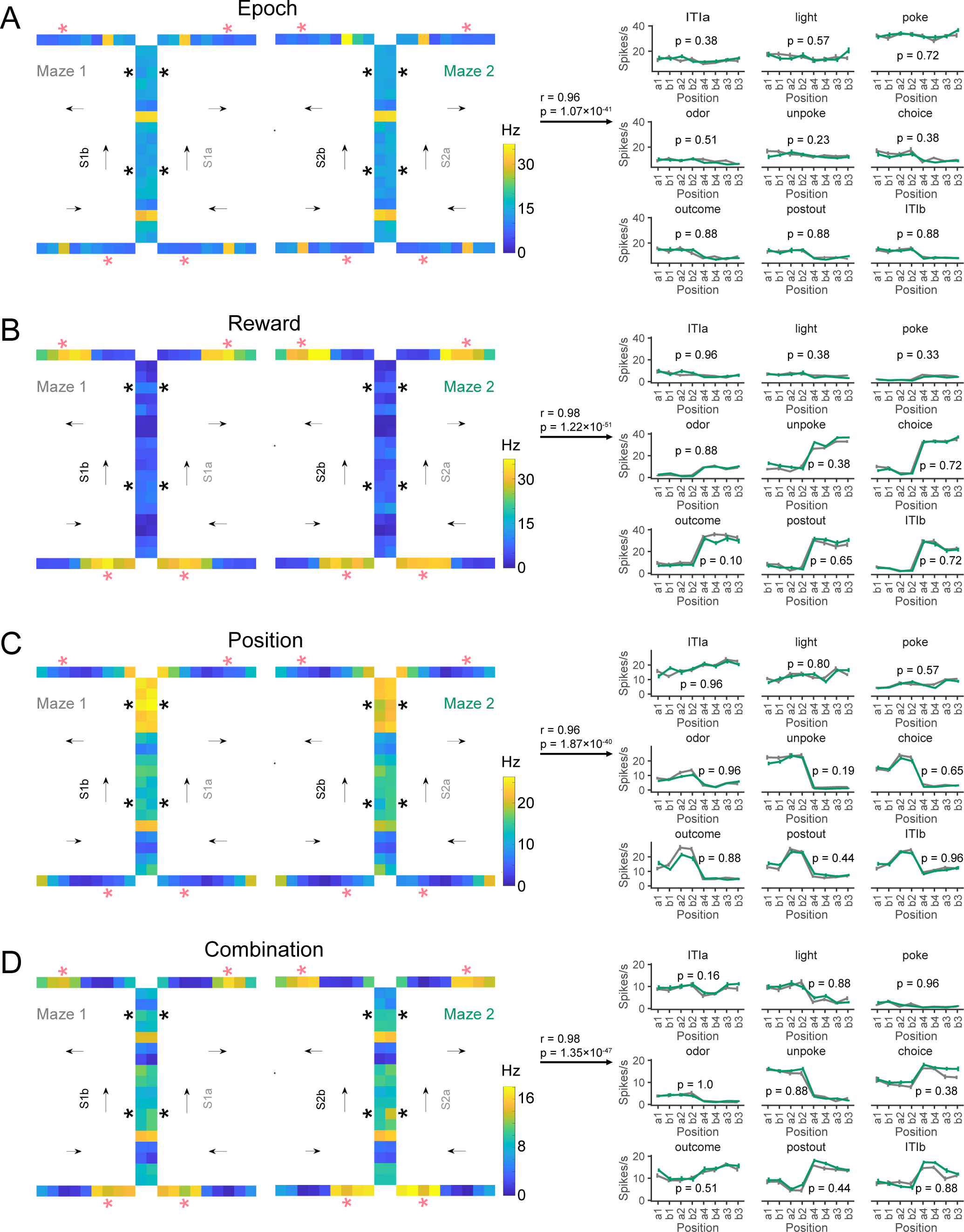
Exemplar units illustrating the generalization of epoch, reward, and positional information across mazes. (A-D) Heatmaps (left) and mean firing rate at each position and epoch (right) for OFC neurons showing generalization of activity related to epoch (A), reward (B), and position (C) or a combination of factors (D). Heatmaps plot activity as described in Figure 2. Line plots show the average firing rate in each epoch at each position in each maze, ordered according to reward on the current and next trials. The gray line represents maze1, and the green line represents maze2. The firing rates were identical between maze1 and maze2 at all epochs in each example (p’s > 0.10, two-sided Wilcoxon rank-sum test). Error bars are SEM. See also Figure S1-3.

### Ventral subiculum inactivation does not affect content or prevalence of schema cells in OFC on an established problem

To quantify the various patterns observed in the single-unit correlates, we designed an algorithmic set of correlational analyses to categorize each unit as representing trial structure, reward, or position and to assess the generalization of that information across mazes. For each unit, we calculated the actual mean firing across correct trials in each of the 9 epochs at each of the 8 positions in the two mazes. We then used correlation coefficients on the resultant pair of 72 x 1 data arrays to determine the generalizability of activity between the two mazes, defining as a schema cell any unit that exhibited a very strong significant correlation (r>0.8 & p<0.01) ^36^. We also defined each unit as influenced by epoch, reward, or position if the correlation across mazes was significant and survived shuffling of information in the other two dimensions (Figure S4). This approach allowed us to classify the activity of each single unit as influenced by each type of information independently, so that we could assess the relationship between them, their generalization, and any effects of time and inactivation.

The analysis identified generalized representations of the two mazes in 47.3% of the units recorded in control sessions (Figure 4A, left). This proportion was similar to the 49.6% identified in neurons recorded in the same rats when ventral subiculum was inactivated in other sessions (Figure 4A, right). While this specific value was based on a somewhat arbitrary criterion, the lack of any effect of inactivation was true regardless of the precise threshold (Figure 4B). Additionally, the proportion of cells that met our criterion was remarkably stable across sessions and repeated inactivation (Figure 4C).

**Figure 4.**
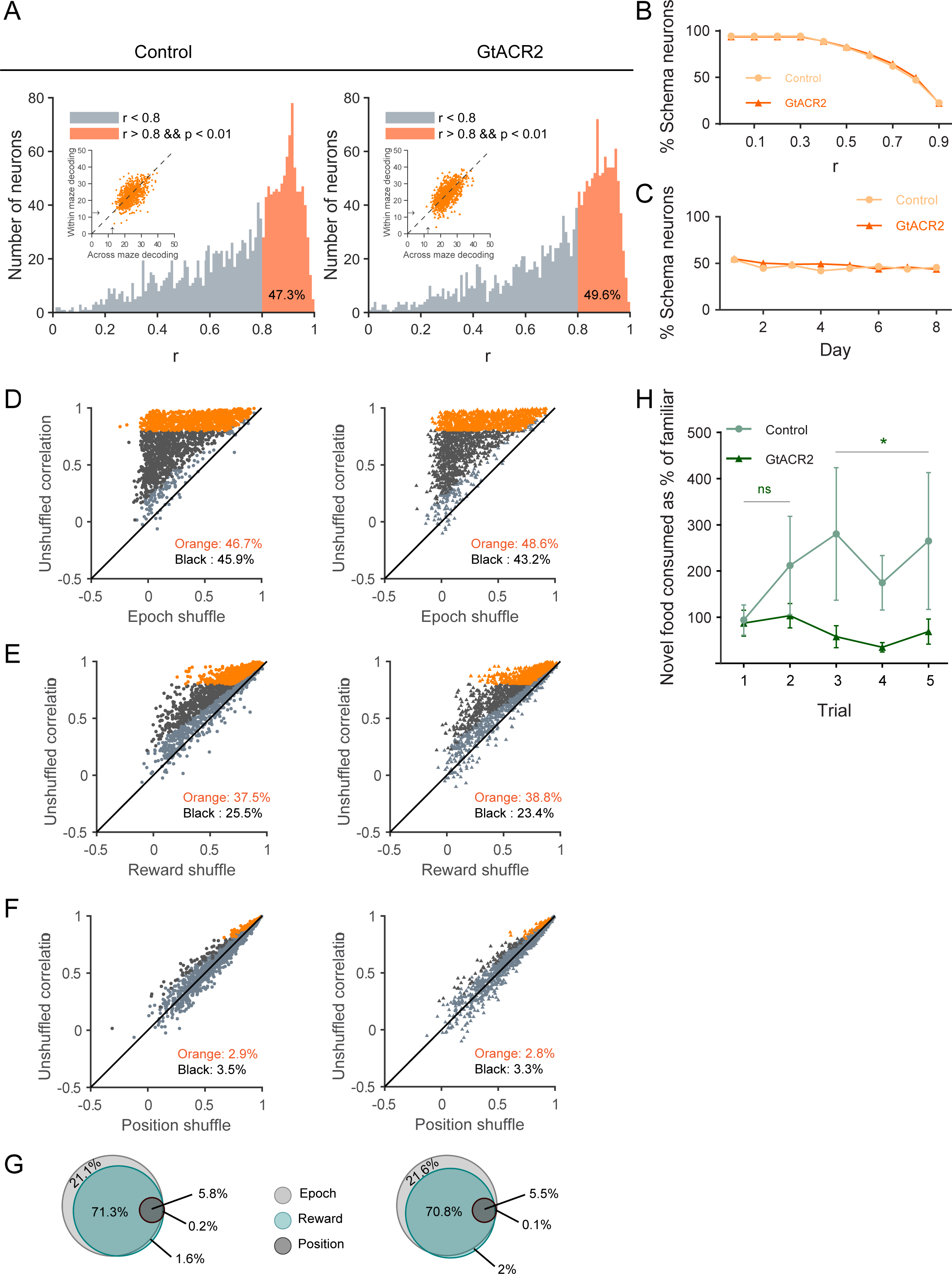
Ventral subiculum inactivation does not affect the content or prevalence of schema cells in OFC on an established problem. (A) Correlation in firing across mazes for all OFC neurons recorded in control and GtACR2 sessions; plot shows the distribution of r scores with the neurons that met the arbitrary cutoff for classification as schema cells (r > 0.8 & p < 0.01) shown in orange. Inset plots: accuracy of decoding position by individual schema neurons, where → denotes chance decoding of 12.5%. One-way ANOVA showed accuracy was similar for decoding within and across mazes for neurons in control (F_3,711_ = 0.068; p = 0.79; η_p_ = 1.8 × 10;) and GtACR2 sessions (F_3,667_= 0.016; p = 0.90; η_p_ = 4.4 × 10), and there was no significant effect of inactivation (within: F_3,689_= 0.45; p = 0.50; η ^2^ = 1.2 × 10^-4^; across: F = 0.35; p = 0.55; η ^2^ = 9.5 × 10^-5^). (B) Percentage of schema neurons at different thresholds for categorization; there was no difference between the two groups in the proportion of neurons at any threshold value (χ^2^ = 0.67; p = 0.41; Chi-squared test). (C) Percentage of schema neurons from control and GtACR2 sessions on each day of training at 0.8 threshold; there was no difference between the two groups in the proportion of neurons at any day (χ^2^’s < 2.3; p’s > 0.86; Chi-squared test). (D-F) Scatter plots show the correlation coefficients of each neuron from the control (left panels) and GtACR2 (right panels) sessions. Y-axes plot the correlation coefficients from unshuffled data; the x-axes plot the mean correlation coefficients obtained after shuffling data (1000x) to disrupt contributions of information related to the epoch (D), reward (E), or position (F). Orange/black cells had actual correlation coefficients larger than 99% of the shuffled results, indicating a significant contribution of the shuffled type of information to the correlated firing patterns. These populations, the size noted on the panels, were not affected by inactivation (χ^2^’s < 2.8; p’s > 0.096; Chi-squared test). (G) Venn diagrams summarize data from panels d-f, showing the fraction of schema neurons recorded in control and GtACR2 sessions that were affected by the shuffling of information related to epoch (light gray), reward (light green), and position (dark gray). Sizes of circles are normalized to the total number of neurons recorded in each group; proportions in each category and overlap between categories were not affected by inactivation (χ^2^’s < 0.40; p’s > 0.54; Chi-squared test). (H) Food consumption across trials in the neophobia task. Lines show novel food consumed per trial as a percentage of familiar food. Light green, control; deep green, GtACR2. A three-way ANOVA revealed a significant main effect of novelty (F_105_= 9.11; p = 0.0032; η ^2^ = 0.081) and a significant interaction between novelty and group (F_105_= 4.05; p = 0.047; η ^2^ = 0.038). Further testing showed a significant difference between groups on the last 3 (F_35_= 7.21; p = 0.01; η ^2^ = 5.6 × 10^-3^) but not the initial 2 trials (F_23_= 0.95; p = 0.34; η ^2^ = 0.042). Error bars are SEM. See also Figure S4 and S6.

Schema cells identified in this manner also maintained relatively high and unchanged classification performance within versus across the two mazes. To show this, we used activity across epochs at each position in one maze as the training set for classification of trials drawn at random from either that maze or the other maze on which that neuron was characterized in a particular session. Using this approach, activity from individual OFC neurons correctly classified the position of test trials ∼25% of the time; this performance did not depend on whether the test trial came from the same or opposite maze as the training set data, and again there was no effect of inactivation (Figure 4A, insets).

Inactivation of ventral subiculum also had very little effect on the content of the generalized representations in OFC. The fractions of units whose correlated activity reflected trial epoch (Figure 4D), reward (Figure 4E), or position (Figure 4F) were entirely unaffected by inactivation, as were the proportion of neurons in each of these categories that met criteria for being schema cells (orange fraction in Figures 4D-F and Figure 4G, see also Figure S6). Thus, inactivation of hippocampal outflow did not dramatically impact established correlates, generalized or not, in the OFC.

Importantly, these negative effects were obtained despite good viral expression and fiber placement (Figure 1D-E). Additionally, we paused the experiment at this point to behaviorally validate the efficacy of inactivation. Rats were trained in a food neophobia task shown to utilize hippocampal processing ^37^. Each day they were given a choice between a novel food pellet and pellets made of their normal chow, and consumption was measured over a 10-minute period, during which the rats received either effective light to inactivate ventral subiculum (n = 3) or ineffective light to serve as control (n = 3). We reasoned that if the appropriate wavelength light were disrupting hippocampal outflow, then the rats receiving it would have difficulty remembering prior exposures to the novel pellet and would show prolonged neophobia relative to their counterparts. Consistent with this prediction, we found that controls increased their consumption of the novel food relative to the familiar whereas the inactivated group did not (Figure 4H). These results provide independent confirmation that light delivery in these rats was acting as expected to disrupt hippocampal output.

### Ventral subiculum inactivation facilitates formation of schema cells in OFC during learning of a new problem

Next, we recorded from OFC during learning of two new problems. The new problems were identical in structure to the first problem (Figure 1B), except that 10 new odors were used for each. Single units were recorded for 10-12 days of training on each problem (9 days of acquisition and then a final day). For this phase, the rats that completed prior training and remained healthy (n = 4) were divided equally into two groups, along with two additional rats which were trained extensively on the initial problem; these rats had approximately similar performance and neural activity during prior training (Figure S5). One group (n = 3) received light to inactivate ventral subiculum during learning of both new problems, while the other (n = 3) received ineffective light to serve as controls. Learning (and the changes in neural correlates) on the two problems was similar, thus we have collapsed them for our analysis.

Rats in both groups quickly learned to discriminate rewarded positions accurately (Figure 5A) and to show differences in their trial initiation latencies reflective of the odor sequences (Figure 5B); these behaviors developed within the first several sessions, which was quicker than the weeks of training required on the initial problem (prior to recording). This is consistent with the development of a schema for learning the basic odor discriminations and the embedded sequence. However, there were differences between the two groups during acquisition of these new problems, which were not evident on the established problem; inactivated rats were faster to stop responding at non-rewarded positions (Figure 5A, scatters, non-rewarded (-)) and failed to distinguish between these positions in their trial initiation speeds (Figure 5B, scatters, P1 vs P2).

**Figure 5.**
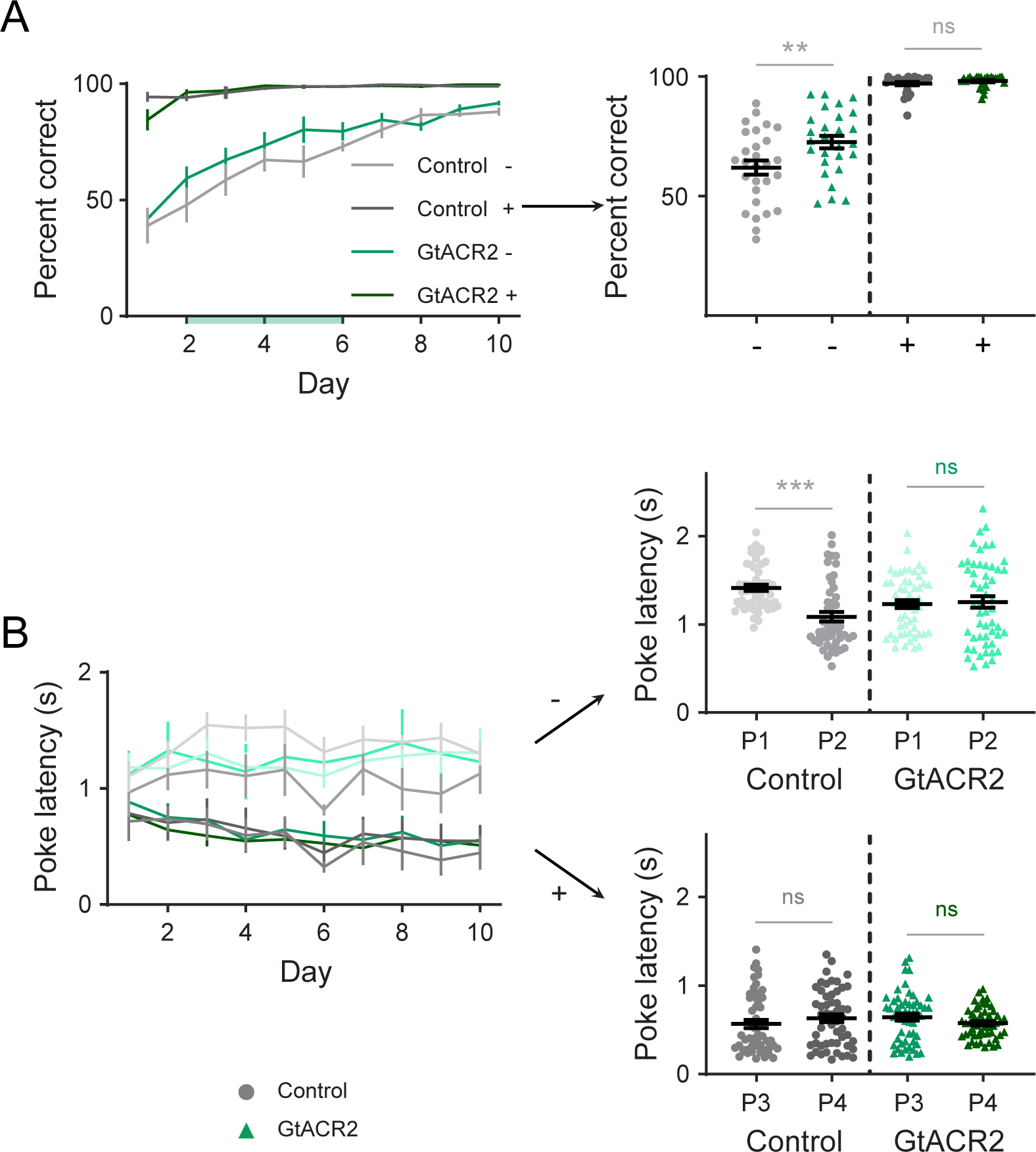
Ventral subiculum inactivation effects on behavior during learning of a new problem. (A-B) Percent correct (A) and trial initiation latencies (B) across days of learning for rats in the control and GtACR2 groups. ANOVAs revealed significant effects of session, trial type, group, an interaction between session and trial type, and an interaction between trial type and group (F’s > 5.4; p’s < 0.021; η_p_^2^’s > 0.03) in the percent correct, reflecting quicker development of the no-go response on non-rewarded positions in the inactivated group at early stages of learning(day 2 to 6) (-in scatter plots: t_53_ = 2.7; p = 9.4 × 10^-3^; Two-tailed test), and a significant main effect of trial type and an interaction between group and trial type (F’s > 5.8, p’s < 0.0007, η ^2^’s > 0.043) in the trial initiation latencies, reflecting a failure of rats in the inactivated group to distinguish the two non-rewarded positions (P1 vs P2 in scatter plots)(Control: t_104_ = 5.2, p = 1.2 × 10^-6^; GtACR2: t_104_ = 0.29, p = 0.77; Two-tailed test). -, non-rewarded trials; +, rewarded trials. Error bars are SEM.***p < 0.001; **p < 0.01; ns, not significant.

Against this backdrop, we recorded single units from each group on each day of training (n = 79-209). Using the approach applied above, we again tracked the development of neural correlates related to trial epoch, reward, and position and their generalization across the two mazes during learning. This analysis showed that in both groups, generalized coding across mazes declined significantly on the new problems in day 1 of training from what had been observed on the established problem (Figure 6A, day 1). From this low, the prevalence of schema cells increased with training, however the increase was significantly greater in rats in which ventral subiculum was inactivated during each trial, such that by day 3, the prevalence in the inactivated group was significantly higher than that in controls and remained higher for the rest of training (Figures 6A-B). The greater prevalence of schema cells in the inactivated group was also evident in the average single-unit positional decoding, which showed a similar decline from prior levels on the established problem in both groups at the start of training and then increased significantly more in the inactivated group than in controls (Figure 6C). Additionally, cells that categorized as schema cells exhibited significantly stronger positional decoding in inactivated rats than in controls (Figure 6D). Thus, inactivation increased both the prevalence and the strength of generalized representations on OFC.

**Figure 6.**
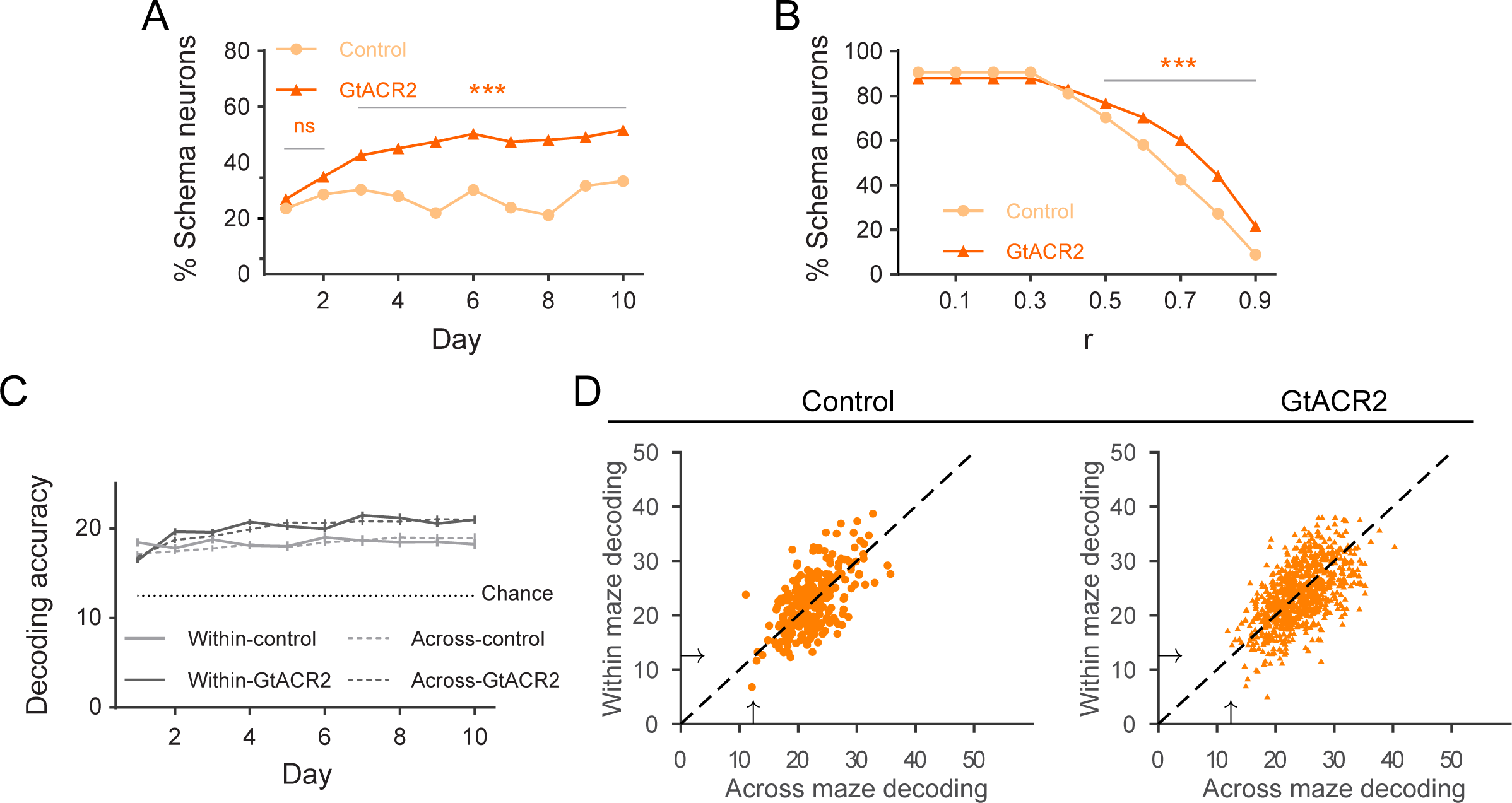
Ventral subiculum inactivation facilitates the formation of schema cells in OFC during learning of a new problem. (A) Percentage of cells recorded on each day that met criterion for schema neurons in each group. The two groups were similar initially and then diverged thereafter (Overall: χ^2^ = 78.9, p = 6.6 x 10^-19^; days 1-2: χ = 1.6; p = 0.21; days 3-10: χ = 83.0, p = 8.0 x 10; Chi-squared test). (B) Percentage of schema neurons in the two groups at different thresholds for categorization. The two groups were similar at low thresholds and then diverged thereafter (Overall: ^2^ = 99.1, p = 2.4 x 10^-23^; r < χ = 9.4, p = 0.002; r >= 0.5: χ = 227.2, p = 2.4 x 10; Chi-squared test). (C) Average single cell decoding of position for all neurons across days. Light gray, control; dark gray, GtACR2. Solid lines, Within the maze; dashed lines, across the maze. Three-way ANOVA revealed a significant main effect of group (F_10_ = 126.1; p = 1.4 × 10^-6^; η ^2^ = 0.93), day (F = 12.4; p = 4.5 × 10^-4^; η ^2^ = 0.93) and a significant interaction between group and day (F_18_= 5.4; p = 9.9 × 10^-3^; η ^2^ = 0.84); there were no main effects or interactions comparing within/across mazes (F’s < 0.79; p’s > 0.40; η ^2^’s < 0.42). (D) Average accuracy of decoding position by individual schema neurons, where → denotes chance decoding of 12.5%. One-way ANOVA showed significant difference of decoding accuracy between the two groups (F _2245_ = 43.4; p = 5.6 × 10^-11^; η ^2^ = 0.019). Error bars are SEM.***p < 0.001; ns, not significant. See also Figure S5.

Notably, though inactivation greatly facilitated the development of generalized representations in OFC, it did not dramatically affect the content of those representations (Figure 7A-D, Figure S7 and Figure S8). The influence of trial epoch remained high throughout training and was not impacted by inactivation (Figures 7A and 7D). The influence of reward and positional information declined in both groups when new problems were introduced (Figures 7B-D), and then increased with training, with a significant increase in the influence of reward (Figures 7B and 7D).

**Figure 7.**
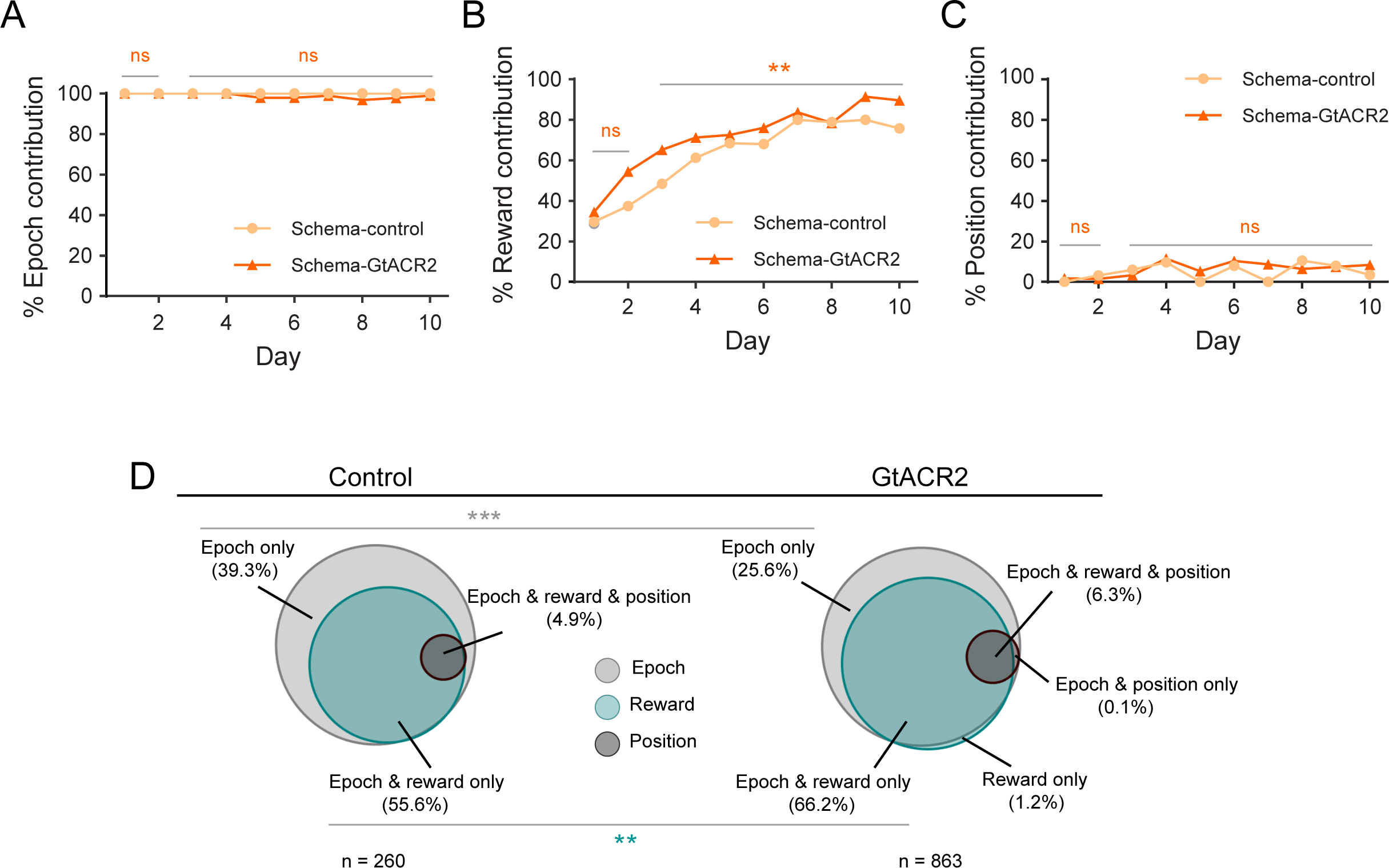
Ventral subiculum inactivation modestly biases the effect of reward on schema cells in OFC during learning of a new problem. (A-C) Percentage of schema neurons whose correlated activity across mazes was affected by shuffling (as in Figure 4D-F) to disrupt information related to epoch (A), reward (B), or position (C). No significant differences between the two groups were observed for either epoch or position (χ^2^’s < 3.3; p’s > 0.067; Chi-squared test), while the influence of reward grew modestly but significantly faster with inactivation (Overall schema: ^2^ = 17.1, p = 3.6 × 10^-5^; days 1-2: ^2^ = 2.2, p = 0.14; days 3-10: ^2^ = 8.9, p = 0.0028; Chi-squared test). (D) Venn diagrams summarize data from panels f-h, showing the fraction of schema neurons recorded in control and GtACR2 sessions that were affected by shuffling of information related to epoch (light gray), reward (light green), and position (dark gray) as in Figure 4G. Sizes of circles are normalized to the total number of neurons recorded in each group, averaged across days (see Figure S8 for same illustration by day); proportions in each category and overlap between categories were affected by inactivation, with an increase in those affected by epoch and reward (χ^2^ = 9.8; p = 0.0018; Chi-squared test) and a corresponding decrease in those affected by epoch only (χ^2^ = 18.4; p = 1.8 × 10^-5^; Chi-squared test). Error bars are SEM.***p < 0.001; **p < 0.01; ns, not significant. See also Figure S7-8.

## Discussion

With the convergence in hypothesized functions of the OFC and HC, it has become increasingly important to test whether and how they coordinate these functions. Nowhere is this more of interest than in the formation and use of generalized cognitive maps or schemas. Here we show that, at least for a single class of problem, “schema cells” are normally hindered or impeded from developing in the OFC by HC output. Notably this was not evident in the expression of an existing schema in a known situation; it was apparent only when an existing schema had to be applied or transferred to a new situation – a new problem pair. Under those conditions, OFC neurons recorded in rats with ventral subiculum inactivated to impair hippocampal output exhibited generalized neural correlates much more quickly and at a much higher prevalence than did neurons recorded in controls, as if hippocampal output were suppressing or interfering with the ability of OFC to port the schema to the new problem. Though surprising, this effect is accords well with evidence of normal learning set formation and even facilitated reversal learning in rats after lesions affecting hippocampal output in rats ^38^. Indeed, even in relatively complex OFC-dependent tasks, such as occasion setting or delayed non-matching, the HC contribution is often limited or non-existent after a problem has been acquired ^16,39^. Enhanced schema cell formation is also consistent with a recent report that hippocampal activity correlates with remapping during new learning in OFC ^28^; inhibiting this interaction during transfer to a new problem might logically have the opposite effect as observed here.

In considering this result, it is important to note that the rats were shaped on the task procedures and an initial problem pair prior to the start of inactivation. Thus while OFC did not require HC to maintain or transfer schema representations to new problems, it is possible that it would depend transiently on HC during this initial stage of learning, as has been shown in other settings ^40^. We also disrupted hippocampal output transiently and at the source rather than lesioning or inactivating a particular subregion within HC or targeting direct projections to or terminals within OFC. We took this approach, effective in prior work ^41^, because we wished to determine how hippocampal outflow affected OFC processing on-the-fly, with minimum time for any compensation, and regardless of the path or other areas involved. Thus, the impact of this sudden inactivation of ventral subiculum on intermediate areas likely shapes our results. One excellent candidate for this might be disruption of interactions with more medial prefrontal areas, which are implicated in switching between or maintaining more context-specific task maps ^32,42-44^, failure of which could contribute to the facilitated generalization observed here.

We also did not attempt to distinguish the contribution of different parts of the hippocampus nor do we argue that inactivation of ventral subiculum completely silences this complex set of areas. We would speculate that processing in dorsal hippocampus is most likely responsible for the interference we observed, since correlates in dorsal hippocampus are strongly related to external information ^45^, whereas recent results have shown ventral hippocampal activity is more strongly influenced by rewards ^46^. Preserved output via pathways like the fornix may also be important in the shape of our effect, especially if these pathways are informationally or functionally biased. In this regard, our results are best viewed as showing one way – rather than the only way - the interaction can go awry.

Our approach also used the chemosensory modality, which may hold a special place for the OFC particularly for rodents, and the goal in this setting – and the information defining the schema - concerns reward, also a feature of the world in which the OFC is often quite interested ^47,48^. While trivial changes – for instance using auditory cues or even spatial locations instead of odors or using food, secondary reinforcers, or even punishment in the place of liquid sucrose reward – seem unlikely to be critical, the finding that HC suppresses schema cell formation in OFC during learning may depend on the type of information that must be generalized to create the schema cells. In our task, schema cells reflect the collapse of information regarding specific features of the positions in the two mazes (i.e. the particular odors or sequences of odors) in favor of information about the rules that predict reward (i.e. the positions in the sequence and/or the reward pattern). The OFC has long been implicated in tracking the conditions predictive of reward ^49-51^, and several studies comparing the encoding in the HC and prefrontal areas – including OFC – have shown that while both represent similar information, representations in the OFC are skewed to reflect the biological significance of the information, while in HC, this influence is much less and instead representations seem to be more attuned to external sensory information ^10,11,23-27,52^. One way to view this is that both OFC and HC contribute to layers of information relevant to cognitive mapping and schemas, with HC focusing on external specifics that define task states and even episodes perhaps, while the OFC warps the map to reflect latent, hidden, or internally defined relevance ^10,53,54^. This predicts that we might see the opposite result if the external cues were the same between two problems, but the rules governing their relationship to reward differed. Under those conditions, inactivation of hippocampal outflow might be predicted to prevent the formation of schema cells in OFC, assuming there were any, since generalization then would depend on the external sensory information. This would be consistent with evidence that HC damage impairs performance in alternation settings and in disambiguating sequences like those used here ^55-57^.

A final aspect of the experiment to consider is that although the results suggest that OFC is not subordinate to HC, it does not comment on the reciprocal relationship. We know that hippocampal activity reflects the influence of many attributes related to reward, hidden states, or goals, information which the OFC is typically tasked with identifying ^19,20,58-61^. Furthermore, prefrontal areas like OFC act to shape neural activity in HC ^62,63^. In settings such as the one used here, we would speculate that the OFC likely influences the HC to compress or generalize where external information differs but internal significance is the same, and to split or distinguish states where external information is the same but internal significance differs ^64^.

Overall, our findings that HC outflow is not necessary to support established schema cells and may at least under some conditions inhibits their emergence during transfer to new problems of a type argue against the idea of a simple feedforward relationship between HC and OFC. Instead, these results strongly favor a model in which OFC and HC operate in parallel and perhaps even somewhat in competition, to extract different features defining cognitive maps and schemas. Within this framework, the OFC is predisposed to form representations that more strongly reflect task relevance and motivational goals, which can be at cross-purposes to the function of HC processing.

### Author Contributions

WZ, JZ, and GS designed the experiments, WZ conducted the behavioral training and single unit recording with assistance from KMC, MPHG and ZZ, and analyzed the data with advice and assistance from JZ and GS. All authors contributed to interpreting the data and writing the manuscript.

## Acknowledgments

The authors thank Dr Ofer Yizhar for his gift of the pAAV-CKIIa-stGtACR2-FusionRed and the histology core at NIDA-IRP, Shiliang Zhang and Dr Marisela Morales for help with preparing and imaging the brains to identify viral expression and fiber and electrode placement. This work was supported by the Intramural Research Program at the National Institute on Drug Abuse and the National Institute on Mental Health. The opinions expressed in this article are the authors’ own and do not reflect the view of the NIH/DHHS. The authors have no conflicts of interest to report.

## Data & Code Availability

The dataset and all scripts used in this study will be made available in an appropriate data archive upon publication.

## Method details

### Subjects

Subjects were four female and four male Long-Evans rats (Charles River Laboratories, 160-300 g) aged ∼3 months at the start of the experiment; analyses revealed no significant main effect nor any interactions with gender in any of the main findings reported in the text, thus rats of different genders were collapsed in all reported data. Rats were housed individually on a 12 h light/dark cycle at the National Institute on Drug Abuse Intramural Research Program (NIDA-IRP). They received ad libitum access to food and water, except during testing periods, during which water was removed from their home cages ∼23h prior to test session. When testing was conducted on consecutive days, they received at least 10 minutes free access to water in their home cages after each test session. All test procedures followed the National Institute of Health guidelines and were approved by the Animal Care and Use Committee at the IRP.

### Surgical procedures

Rats were implanted with drivable bundles of 16 nickel-chromium wires (25-mm diameter; AM Systems) targeting the lateral OFC bilaterally (AP: 3 mm, ML: 3.2 mm). Each wire bundle was housed in a stainless-steel hypodermic tubing (27-gauge, 0.01625-inch OD, 0.01025-inch ID) and cut with a pair of fine bone-cutting spring scissors (16144-13; Fine Science Tools) to extend 1.5 – 2mm beyond the end of the cannula inside the brain. The tips of the wires were initially implanted 4.2mm ventral to the brain surface and subsequently advanced during the retraining period to obtain high quality stable recordings. During the same surgery, pAAV-CKIIa-stGtACR2-FusionRed ^34^ (Addgene viral prep #105669-AAV1) was infused bilaterally in the ventral subiculum (6.5mm posterior and ± 4.5mm lateral of bregma), and optical fibers (Thorlabs) were positioned over each injection site. Rats were given Cephalexin (15 mg/kg) orally twice a day for two weeks to prevent infection after surgery. Rats were allowed ∼4 to 5 weeks to recover from surgery and to allow viral expression before they began reminder training as described below.

### Food neophobia test

Between recording on the initial and learning problems, the rats were food deprived for 48 hours and then underwent testing in a hippocampal-dependent neophobia task to confirm the functional efficacy of GtACR2 at inactivating hippocampal outflow. Consumption was tested across 5 sessions in which rats were placed into a box individually and presented with similar amounts of a familiar (normal chow) and novel food (bacon or banana flavored sucrose pellets) for 10 minutes, while receiving light effective at activating the GtACR2 molecule or ineffective light of a similar power as a control. Food ramekins were situated on either side in front of the rat, and the orientation of the two foods were counterbalanced within and across days. Both the box and ramekins were cleaned with wet hand towels and dried after each rat was tested to reduce any influence of previous tests. Any food remaining after 10 minutes was collected and weighed to determine consumption.

### Dual figure 8 odor sequence task

Training and recording sessions were conducted in aluminum chambers (∼18 inches on a side) outfitted with panels containing an odor port flanked by two fluid-delivery wells, which were monitored by infrared beam sensors across each opening. The odor port was connected to a custom olfactometer, which allowed unique odor cues to be delivered with a rapid onset and in a controlled fashion, and the right well allowed delivery of a sucrose reward, all of which was monitored and controlled by custom behavioral software written in C++. Each trial began with illumination of 2 house lights located above the odor port, which signaled the availability of an odor cue at the port. A stable nose poke (500ms) at the odor port-initiated odor delivery (500ms), after which the rats were free to withdraw from the odor port and make a “go” response at the right fluid well. A response on positive trials was considered correct and led to the delivery of 90μl of a 5% sucrose solution after a random delay (400-1,500ms), and extinction of the house lights on well exit to start the ITI; a response on negative trials was considered an error and led to immediate extinction of the house lights. If no response was made, which generally only occurred on negative trials where it was correct, the house lights were extinguished after a 2s period, and the trial was considered a “no-go”. The ITI period was 4s after correct trials and 6s after errors, beginning when the house lights went off.

One of 10 odors was delivered to the odor port on each trial, and the ten odors were organized into two fixed sequences, the pattern of which constituted what we refer to as non-spatial figure-8 mazes (maze 1 and maze 2). Each maze can be broken down into two subsequences as illustrated below, with numbers indicating the unique odor cue and positive (+) and negative (−) symbols to indicate reward availability:

Maze 1:

S1a: 0-, 1−, 2+, 2+;

S1b: 0-, 1−, 3+, 4+;

Maze 2:

S2a: 5-, 6−, 7+, 7+;

S2b: 5-, 6−, 8+, 9+;

Each daily training session consisted of 320 trials, divided into 4 blocks of 80 trials involving one of the two mazes. Blocks were presented randomly in one of the two orders: maze1, maze2, maze1, maze2 or maze2, maze1, maze2, maze1. Before recording, rats were shaped to nosepoke at the odor port and then respond at the well for a reward, after which they were immediately introduced to the odor sequences. All rats (n = 8) were trained until they were able to reliably complete the 320 trials each day at a criterion of >75% correct performance on every position (35-45 sessions), after which electrode arrays were implanted in the OFC. Subsequently, all rats (n = 8) received additional reminder training after surgery, after which recording began.

Recording during accurate performance on the initial maze pair was divided into control and inactivation sessions. As described in the optogenetic methods, light with a wavelength effective at activating the GtACR2 molecule was delivered during inactivation sessions, whereas light of a similar power but ineffective wavelength was delivered during control sessions. All rats that participated in this part of the study (n = 6) underwent both conditions, with the session type alternating randomly except that the same condition could not occur on 3 consecutive days. This resulted in 8 control and 8 inactivation sessions from all rats but one, which fell ill after completing only 3 sessions of each type. Data from these sessions are included but effects reported do not depend upon it. An additional rat from the original 6 grew ill in the weeks following the end of recording and had to be removed from the remainder of the study.

After recording on the initial maze pair, the 4 rats that remained, along with 2 new rats, were divided into control and inactivation groups (n’s = 3). The two new rats - one in each group – were required to replace the rats that were removed during prior training. These new rats received shaping, surgery, recovery, and additional reminder training on the initial problem as described above to parallel the training of the original rats. Critically, the original rats placed in the two groups had exhibited similar proportions of schema cells during the prior training, and the new rats that were added exhibited proportions of schema cells during their reminder training like the original rats (Figure S5). Single-unit activity was recorded as the rats in these two groups learned two new maze pairs. Each new maze pair consisted of 10 new odor cues, arranged in sequences with the same structure as the initial maze pair; recording continued for 10-12 days on each new pair, and the resultant analyses focused on days 1-9 and the final day of recording.

### Single-unit recording

Plexon OmniPlex (Plexon, Dallas, TX) systems were used to record electrophysiological signals. Neural signals were digitized, amplified, and bandpass filtered (250 – 8,000Hz) to isolate spike activity, and a threshold was set manually for each active channel to capture unsorted spikes. Timestamps for behavioral events were sent to the Plexon system, synchronized, and recorded alongside the neural activity. Spikes were sorted to remove noise and identify single units offline using Offline Sorter (v.4.0; Plexon) with a built-in template-matching algorithm. Sorted files were sent to NeuroExplorer (Nex Technologies, Colorado Springs, CO) to extract unit and behavioral event timestamps, which were then exported as MATLAB (2021b; MathWorks) formatted files for further analysis. Electrodes were not advanced within a given problem; however we make no claims regarding whether single units recorded on different days within the same problem are the same or different neurons. The electrodes were advanced ∼120 μm to change the neural population being sampled between odor problems.

### Optogenetic stimulation

Light was delivered using a combined optogenetic/electrical commutator interfaced with custom-made 1.25 mm FC ferrules (Thorlabs). To inactivate ventral subiculum, 465 nm (6–10 mW power output) light was delivered to activate GtACR2 and suppress neural activity. As a control, 630 nm light (6–10 mW power output) was delivered. This wavelength falls outside the frequency sensitivity range of GtACR2 molecule ^35^. In the figure 8 task, light was delivered continuously (i.e., not pulsed) during each trial, starting with house light illumination, and terminating after outcome delivery. During recording on the initial maze, each rat received both control and inactivating wavelengths of light in different sessions, alternating pseudorandomly; during recording on subsequent mazes, each rat received either control or inactivating light. In the neophobia task, light was delivered continuously for the full 10-minute test period, and each rat received either control or inactivating light.

### Peri-event time epochs

Each trial was divided into nine epochs associated with different events:

1) ITIa = time from 500ms before to light onset

2) light = 0ms before to 500ms after light onset

3) poke = 0ms before to 500ms after odor port nosepoke

4) odor = 0ms before to 500ms after odor delivery

5) unpoke = 0ms before to 500ms after odor port unpoke

6) choice = 0ms before to 500ms after well entry (or trial termination)

7) outcome = 0ms before to 500ms after reward delivery (or 500ms after trial termination)

8) postout = 500ms after end of outcome period

9) ITIb = 500ms after end of postout period

The spike trains for every isolated unit were aligned with the onset of each event. Spike number was counted with a bin = 100 ms. A Gaussian kernel (s = 50 ms) was used to smooth the spike train on each trial.

### Quantification and statistical analyses

The number of rats and neurons were not predetermined by any statistical methods but are comparable to those reported in previous publications from our and other labs. All data were analyzed using MATLAB (R2021b; MathWorks). Error bars in figures denote the standard error of the mean.

### Single neuron selectivity and percent of maximal firing analysis

The firing rate of each neuron was assessed by three-way ANOVA to determine whether it was selective to epoch, position, or value (p<0.01). For each epoch-selective neuron, the maximal firing rate at all epochs was found, then the percentage of each epoch was calculated. For each position-selective neuron, the maximal firing rate across all positions (P1, P2, P3, P4) was found, and the respective percentage for each position was calculated. For reward-selective neurons, the maximal firing rate at rewarded or non-rewarded trials was assessed, and the percentage for each reward category was calculated.

### Calculation of spatial correlations

We performed correlation analysis to check the activity of data sets from one maze to the other. Pairwise Pearson correlation was calculated by applying the corrcoef function of Matlab on all cells for mean firing rate of each trial type of all correct trials binned with 100 ms resolution. The eight trial types and nine epochs resulted in a 72 x 1 matrix for each maze. We determine schema cells as the ones with strong correlation (correlation coefficient > 0.8) when p <0.01. These original correlation coefficients (r1) will be used to determine the following analysis’s position-specific, epoch-specific, position/value-specific and value-specific contributions.

Epoch-specific: disrupt schema correlates that are epoch-specific but not ones that are purely value or position specific by shuffle data between epochs within each maze but keeping reward/position category the same 1000 times. Then determine the correlation coefficient for the two mazes using the corrcoef function of Matlab again. We determine a neuron as epoch-specific when r1 was more than 99% of the shuffled correlation coeffects using prctile function of Matlab.

Reward specific: disrupt schema correlates that are reward/position specific but not ones that are purely epoch specific by shuffle data between rewarded/nonrewarded positions within each maze but keeping epoch the same 1000 times. Then determine the correlation coefficient for the two mazes using the corrcoef function of Matlab again. We determine a neuron as reward/position specific when r1 was more than 99% of the shuffled correlation coeffects using prctile function of Matlab.

Position-specific: disrupt schema correlates that are position-specific but not ones that are purely epoch or reward specific by shuffling data between positions in each maze but keeping reward category and epoch the same 1000 times. Then determine the correlation coefficient for the two mazes using the corrcoef function of Matlab again. We determine a neuron as position-specific when r1 was more than 99% of the shuffled correlation coeffects using prctile function of Matlab.

### Single-cell decoding

All 20 trials for each trial type were included for 9 epochs, resulting in a 160 (no. of trials) × 9 (no. of epochs) matrix. Then, randomly leave one trial of each 8 trial types out to get an 8 × 9 test set from maze1(within maze), while the rest 152 × 9 matrix was used to train the model. The same size of matrix (8 × 9) from maze 2 with an identical trial index as the test set from maze 1 was used for cross-maze decoding of trial types. Based on the matrix of firing rate, a linear multi-class classification (LIBLINEAR ^65^, https://www.csie.ntu.edu.tw/~cjlin/liblinear/) was trained to classify 8 trial types. This procedure was repeated 1000 times, then the average decoding accuracy within the maze and across the maze was calculated for each cell. The chance level was 1/8 for each cell.

## References

1. Tolman, E.C. (1948). Cognitive maps in rats and men. Psychological Review 55, 189–208.

2. Behrens, T.E., Muller, T.H., Whittington, J.C.R., Mark, S., Baram, A.B., Stachenfeld, K.L., and Kurth-Nelson, Z. (2018). What is a cognitive map? Organizing knowledge for flexible behavior. Neuron 100, 490–509.

3. O’Keefe, J., and Nadel, L. (1978). The Hippocampus as a Cognitive Map (Clarendon Press).

4. Wilson, R.C., Takahashi, Y.K., Schoenbaum, G., and Niv, Y. (2014). Orbitofrontal cortex as a cognitive map of task space. Neuron 81, 267–279.

5. Eichenbaum, H. (2000). Hippocampus: Mapping or memory? Current Biology 10, R785–R787.

6. Gershman, S.J., and Niv, Y. (2010). Learning latent structure: carving nature at its joints. Current Opinion in Neurobiology 20, 251–256.

7. Niv, Y. (2019). Learning task-state representations. Nature Neuroscience 22, 1544–1553.

8. Zhou, J., Jia, C., Montesinos-Cartegena, M., Gardner, M.P.H., Zong, W., and Schoenbaum, G. (2021). Evolving schema representations in orbitofrontal ensembles during learning. Nature 590, 606–611.

9. Baraduc, P., Duhamel, J.-R., and Wirth, S. (2019). Schema cells in the macaque hippocampus. Science 363, 635–639.

10. Farovik, A., Place, R.J., McKenzie, S., Porter, B., Munro, C.E., and Eichenbaum, H. (2015). Orbitofrontal cortex encodes memories within value-based schemas and represents contexts that guide memory retrieval. Journal of Neuroscience 35, 8333–8344.

11. McKenzie, S., Frank, A.J., Kinsky, N.R., Porter, B., Riviere, P.D., and Eichenbaum, H. (2014). Hippocampal representation of related and opposing memories develop with distinct, heirarchically organized neural schemas. Neuron 83, 202–215.

12. Lipton, P.A., Alvarez, P., and Eichenbaum, H. (1999). Crossmodal associative memory representations in rodent orbitofrontal cortex. Neuron 22, 349–359.

13. Wikenheiser, A.M., Gardner, M.P.H., Mueller, L.E., and Schoenbaum, G. (2021). Spatial representations in the rat orbitofrontal cortex. Journal of Neuroscience 41, 6933–6945.

14. Young, J.J., and Shapiro, M.L. (2011). Dynamic coding of goal-directed paths by orbital prefrontal cortex. Journal of Neuroscience 31, 5989–6000.

15. Constantinescu, A.O., O’Reilly, J.X., and Behrens, T.E. (2016). Organizing conceptual knowledge in humans with a gridlike code. Science 352, 1464–1468.

16. Otto, T., and Eichenbaum, H. (1992). Complementary roles of the orbital prefrontal cortex and the perirhinal-entorhinal cortices in an odor-guided delayed-nonmatching-to-sample task. Behavioral Neuroscience 106, 762–775.

17. Bradfield, L.A., Leung, B.K., Boldt, S., Liang, S., and Balleine, B.W. (2020). Goal-directed actions transiently depend on dorsal hippocampus. Nature Neuroscience 23, 1194–1197.

18. Miller, K.J., Botvinick, M.M., and Brody, C.D. (2017). Dorsal hippocampus contributes to model-based planning. Nature Neuroscience 20, 1269–1276.

19. Wikenheiser, A.M., and Redish, A.D. (2015). Hippocampal theta sequences reflect current goals. Nature Neuroscience 18, 289–294.

20. Knudsen, E.B., and Wallis, J.D. (2021). Hippocampal neurons construct a map of an abstract value space. Cell 184, 4640–4650.

21. Vikbladh, O.M., Meager, M.R., King, J., Blackmon, K., Devinsky, O., Shohamy, D., Burgess, N., and Daw, N.D. (2019). Hippocampal contributions to model-based planning and spatial memory. Neuron 102, 683–693.

22. Wimmer, G.E., and Shohamy, D. (2012). Preference by association: How memory mechanisms in the hippocampus bias decisions. Science 338, 270–273.

23. Young, B.J., Otto, T., Fox, G.D., and Eichenbaum, H. (1997). Memory representation within the parahippocampal region. Journal of Neuroscience 17, 5183–5195.

24. Ramus, S.J., and Eichenbaum, H. (2000). Neural correlates of olfactory recognition memory in the rat orbitofrontal cortex. Journal of Neuroscience 20, 8199–8208.

25. Zhou, J., Montesinos-Cartegena, M., Wikenheiser, A.M., Gardner, M.P.H., Niv, Y., and Schoenbaum, G. (2019). Complementary task structure representations in hippocampus and orbitofrontal cortex during an odor sequence task. Current Biology 29, 3402–3409.

26. Place, R., Farovik, A., Brockmann, M., and Eichenbaum, H. (2016). Bidirectional prefrontal-hippocampal interactions support context-guided memory. Nature Neuroscience 19, 992–994.

27. Samborska, V., Butler, J.L., Walton, M.E., Behrens, T.E.J., and Akam, T. (2022). Complementary task representations in hippocampus and prefrontal cortex for generalising the structure of problems. Nature Neuroscience 25, 1314–1326.

28. Riceberg, J.S., Srinivasan, A., Guise, K.G., and Shapiro, M.L. (2022). Hippocampal signals modify orbitofrontal representations to learn new paths. Current Biology 32, 3407–3413.

29. Nardin, M., Kaefer, K., and Csicsvari, J. (2021). The generalized spatial representation in the prefrontal corterx is inherited from the hippocampus. BioRxiv. https://doi.org/10.1101/2021.09.30.462269.

30. Schuck, N.W., and Niv, Y. (2019). Sequential replay of nonspatial task states in human hippocampus. Science 364, 6447.

31. Mizrak, E., Bouffard, N.R., Libby, L.A., Boorman, E.D., and Ranganath, C. (2021). The hippocampus and orbitofrontal cortex jointly represent task structure during memory-guided decision making. Cell Reports 37, 110065.

32. Preston, A.R., and Eichenbaum, H. (2013). Interplay of hippocampus and prefrontal cortex in memory. Current Biology 23, R764–773.

33. Whittington, J.C.R., McCaffary, D., Bakermans, J.J.W., and Behrens, T.E.J. (2022). How to build a cognitive map. Nature Neuroscience 25, 1257–1272.

34. Mahn, M., Gibor, L., Patil, P., Cohen-Kashi, M.K., Oring, S., Printz, Y., Levy, R., Lampl, I., and Yizhar, O. (2018). High-efficiency optogenetic silencing with soma-targeted anion-conducting channelrhodopsins. Nature Communications 9, 4125.

35. Govorunova, E.G., Sineshchekov, O.A., Janz, R., Liu, X., and Spudich, J.L. (2015). Natural light-gated anion channels: a family of microbial rhodopsins for advanced optogenetics. Science 349, 647–650.

36. Chan, Y.H. (2003). Biostatistics 104: correlational analysis. Singapore Med J 44, 614–619.

37. Kelley, A.E. (1999). Neural integrative activities of nucleus accumbens subregions in relation to learning and memory. Psychobiology 27, 198–213.

38. Eichenbaum, H., Fagan, A., and Cohen, N.J. (1986). Normal olfactory discrimination learning set and facilitation of reversal learning after medial-temporal damage in rats: Implications for an account of preserved learning abilities in amnesia. Journal of Neuroscience 6, 1876–1884.

39. Fraser, K.M., and Janak, P.H. (2022). Basolateral amygdala and orbitofrontal cortex, but not dorsal hippocampus, are necessary for the control of reward-seeking by occassion setters. Psychopharmacology 240, 623–635.

40. Tse, D., Langston, R.F., Kakeyama, M., Bethus, I., Spooner, P.A., Wood, E.R., Witter, M.P., and Morris, R.G.M. (2007). Schemas and memory consolidation. Science 316, 76–82.

41. Wikenheiser, A.M., Marrero-Garcia, Y., and Schoenbaum, G. (2017). Suppression of ventral hippocampal output impairs integrated orbitofrontal encoding of task structure. Neuron 95, 1197–1207.

42. Sharpe, M.J., Stalnaker, T.A., Schuck, N.W., Killcross, A.S., Schoenbaum, G., and Niv, Y. (2018). An integrated model of action selection: distinct modes of cortical control of striatal decision making. Annual Review of Psychology 70, 53–76.

43. Eichenbaum, H. (2017). Prefrontal-hippocampal interactions in episodic memory. Nature Reviews Neuroscience 18, 547–558.

44. Alonso, A., Meij, v.d., Tse, D., and Genzel, L. (2020). Naive to expert: considering the role of previous knowledge in memory. Bain and Neuroscience Advances 4, 1–17.

45. Jung, M.W., Weiner, S.I., and McNaughton, B.L. (1994). Comparison of spatial firing characteristics of units in dorsal and ventral hippocampus of the rat. Journal of Neuroscience 14, 7347–7356.

46. Lee, I., and Jin, S.-W. (2021). Differential encoding of place value between the dorsal and intermediate hippocampus. Current Biology 31, 3053–3072.

47. Knudsen, E.B., and Wallis, J.D. (2022). Taking stock of value in the orbitofrontal cortex. Nature Reviews Neuroscience 23, 428–438.

48. Padoa-Schioppa, C., and Conen, K.E. (2017). Orbitofrontal cortex: a neural circuit for economic decisions. Neuron 96, 736–754.

49. Thorpe, S.J., Rolls, E.T., and Maddison, S. (1983). The orbitofrontal cortex: neuronal activity in the behaving monkey. Experimental Brain Research 49, 93–115.

50. Gallagher, M., McMahan, R.W., and Schoenbaum, G. (1999). Orbitofrontal cortex and representation of incentive value in associative learning. Journal of Neuroscience 19, 6610–6614.

51. Izquierdo, A.D., Suda, R.K., and Murray, E.A. (2004). Bilateral orbital prefrontal cortex lesions in rhesus monkeys disrupt choices guided by both reward value and reward contingency. Journal of Neuroscience 24, 7540–7548.

52. Ross, R.S., LoPresti, M.L., Schon, K., and Stern, C.E. (2013). Role of the hippocampus and orbitofrontal cortex during the disambiguation of social cues in working memory. Cognitive, Affective, & Behavioral Neuroscience 13, 900–915.

53. Costa, K.M., Scholz, R., Lloyd, K., Moreno-Castilla, P., Gardner, M.P.H., Dayan, P., and Schoenbaum, G. (2023). The role of the lateral orbitofrontal cortex in creating cognitive maps. Nature Neuroscience 26, 107–115.

54. Schuck, N.W., Cai, M.B., Wilson, R.C., and Niv, Y. (2016). Human orbitofrontal cortex represents a cognitive map of state space. Neuron 91, 1402–1412.

55. Agster, K.L., Fortin, N.J., and Eichenbaum, H. (2002). The hippocampus and disambiguation of overlapping sequences. Journal of Neuroscience 22, 5760–5768.

56. Fortin, N.J., Agster, K.L., and Eichenbaum, H. (2002). Critical role of the hippocampus in memory for sequences of events. Nature Neuroscience 5, 458–462.

57. Hock, B.J., and Bunsey, M.D. (1998). Differential effects of dorsal and ventral hippocampal lesions. Journal of Neuroscience 18, 7027–7032.

58. Gauthier, J.L., and Tank, D.W. (2018). A dedicated population for reward coding in the hippocampus. Neuron 99, 179–193.

59. Carey, A.A., Tanaka, Y., and van der Meer, M.A.A. (2019). Reward revaluation biases hippocampal replay content away from the preferred outcome. Nature Neuroscience 22, 1450–1459.

60. Zeithamova, D., Gelman, B.D., Frank, L., and Preston, A.R. (2018). Abstract representation of prospective reward in the hippocampus. Journal of Neuroscience 38, 10093–10101.

61. Wang, F., Schoenbaum, G., and Kahnt, T. (2020). Interactions between human orbitofrontal cortex and hippocampus support model-based inference. PloS Biology 18, e3000578.

62. Guise, K.G., and Shapiro, M.L. (2017). Medial prefrontal cortex reduces memory interference by modifying hippocampal encoding. Neuron 94, 183–192.

63. Garvert, M.M., Saanum, T., Schulz, E., Schuck, N.W., and Doeller, C.F. (2023). Hippocampal spatio-predictive cognitive maps adaptively guide reward generalization. Nature.

64. Duvelle, E., Grieves, R.M., and van der Meer, M.A. (2022). Temporal context and latent state inference in the hippocampal splitter signal. eLIFE 12, e82357.

65. Fan, R.-E., Chang, K.-W., Hsieh, X.-R., and Lin, C.-J. (2008). LIBLINEAR: A library for large linear classification. Journal of Machine Learning Research 9, 1871–1874.

