## Supplemental figures and captions for "Schema cell formation in orbitofrontal cortex is suppressed by hippocampal output"

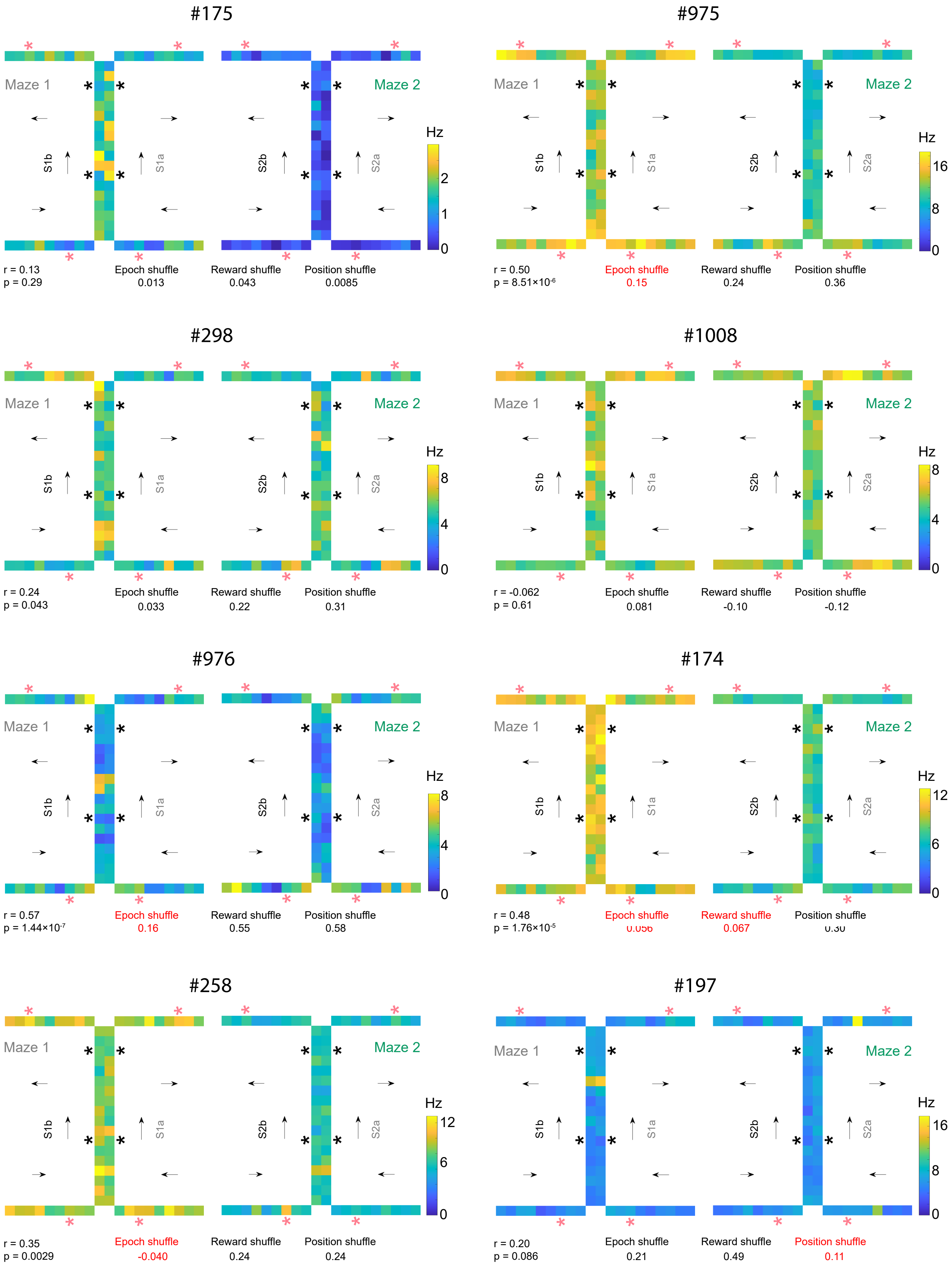

#175
Maze 1
Maze 2
S1b
S1a
S2b
S2a
Hz
0
1
2
r = 0.13
p = 0.29
Epoch shuffle
0.013
Reward shuffle
0.043
Position shuffle
0.0085
#975
Maze 1
Maze 2
S1b
S1a
S2b
S2a
Hz
0
8
16
r = 0.50
p = 8.51×10-6
Epoch shuffle
0.15
Reward shuffle
0.24
Position shuffle
0.36
#298
Maze 1
Maze 2
S1b
S1a
S2b
S2a
Hz
0
4
8
r = 0.24
p = 0.043
Epoch shuffle
0.033
Reward shuffle
0.22
Position shuffle
0.31
#1008
Maze 1
Maze 2
S1b
S1a
S2b
S2a
Hz
0
4
8
r = -0.062
p = 0.61
Epoch shuffle
0.081
Reward shuffle
-0.10
Position shuffle
-0.12
#976
Maze 1
Maze 2
S1b
S1a
S2b
S2a
Hz
0
4
8
r = 0.57
p = 1.44×10-7
Epoch shuffle
0.16
Reward shuffle
0.55
Position shuffle
0.58
#174
Maze 1
Maze 2
S1b
S1a
S2b
S2a
Hz
0
6
12
r = 0.48
p = 1.76×10-5
Epoch shuffle
0.056
Reward shuffle
0.067
Position shuffle
0.30
#258
Maze 1
Maze 2
S1b
S1a
S2b
S2a
Hz
0
6
12
r = 0.35
p = 0.0029
Epoch shuffle
-0.040
Reward shuffle
0.24
Position shuffle
0.24
#197
Maze 1
Maze 2
S1b
S1a
S2b
S2a
Hz
0
8
16
r = 0.20
p = 0.086
Epoch shuffle
0.21
Reward shuffle
0.49
Position shuffle
0.11

**Figure S1. Exemplar units illustrating maze-unique (i.e. non-generalized) firing patterns in OFC.**

A pair of heatmaps are shown for each neuron, plotting average activity in each epoch at each position in each maze. Arrows represent sequence directions. Red * marks the reward epoch on rewarded trial types (P3 and P4), while black * marks the reward epoch for non-rewarded trial types (P1 and P2). The correlation coefficients before and after shuffling of epoch, reward, and position are presented in the lower panel of each figure; numbers shown in red were significantly affected by shuffling.

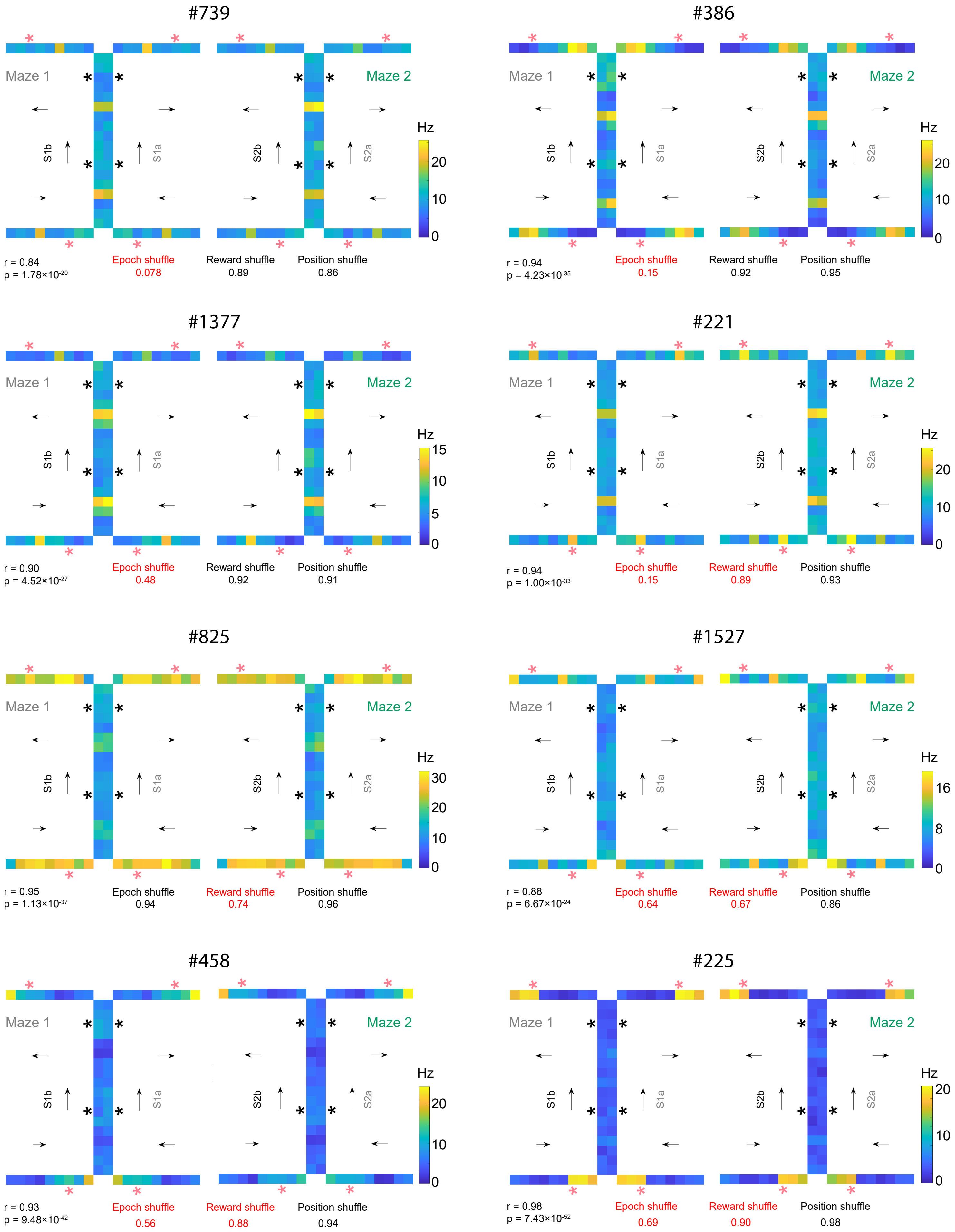
#739
Maze 1
Maze 2
S1b
S1a
S2b
S2a
Hz
r = 0.84
p = 1.78×10-20
Epoch shuffle
0.078
Reward shuffle
0.89
Position shuffle
0.86
#386
r = 0.94
p = 4.23×10-35
Epoch shuffle
0.15
Reward shuffle
0.92
Position shuffle
0.95
#1377
r = 0.90
p = 4.52×10-27
Epoch shuffle
0.48
Reward shuffle
0.92
Position shuffle
0.91
#221
r = 0.94
p = 1.00×10-33
Epoch shuffle
0.15
Reward shuffle
0.89
Position shuffle
0.93
#825
r = 0.95
p = 1.13×10-37
Epoch shuffle
0.94
Reward shuffle
0.74
Position shuffle
0.96
#1527
r = 0.88
p = 6.67×10-24
Epoch shuffle
0.64
Reward shuffle
0.67
Position shuffle
0.86
#458
r = 0.93
p = 9.48×10-42
Epoch shuffle
0.56
Reward shuffle
0.88
Position shuffle
0.94
#225
r = 0.98
p = 7.43×10-52
Epoch shuffle
0.69
Reward shuffle
0.90
Position shuffle
0.98

**Figure S2. Exemplar units illustrate the generalization of epoch and reward information across mazes in OFC.**

A pair of heatmaps are shown for each neuron, plotting average activity in each epoch at each position in each maze. Arrows represent sequence directions. Red * marks the reward epoch on rewarded trial types (P3 and P4), while black * marks the reward epoch for non-rewarded trial types (P1 and P2). The correlation coefficients before and after shuffling of epoch, reward, and position are presented in the lower panel of each figure; numbers shown in red were significantly affected by shuffling.

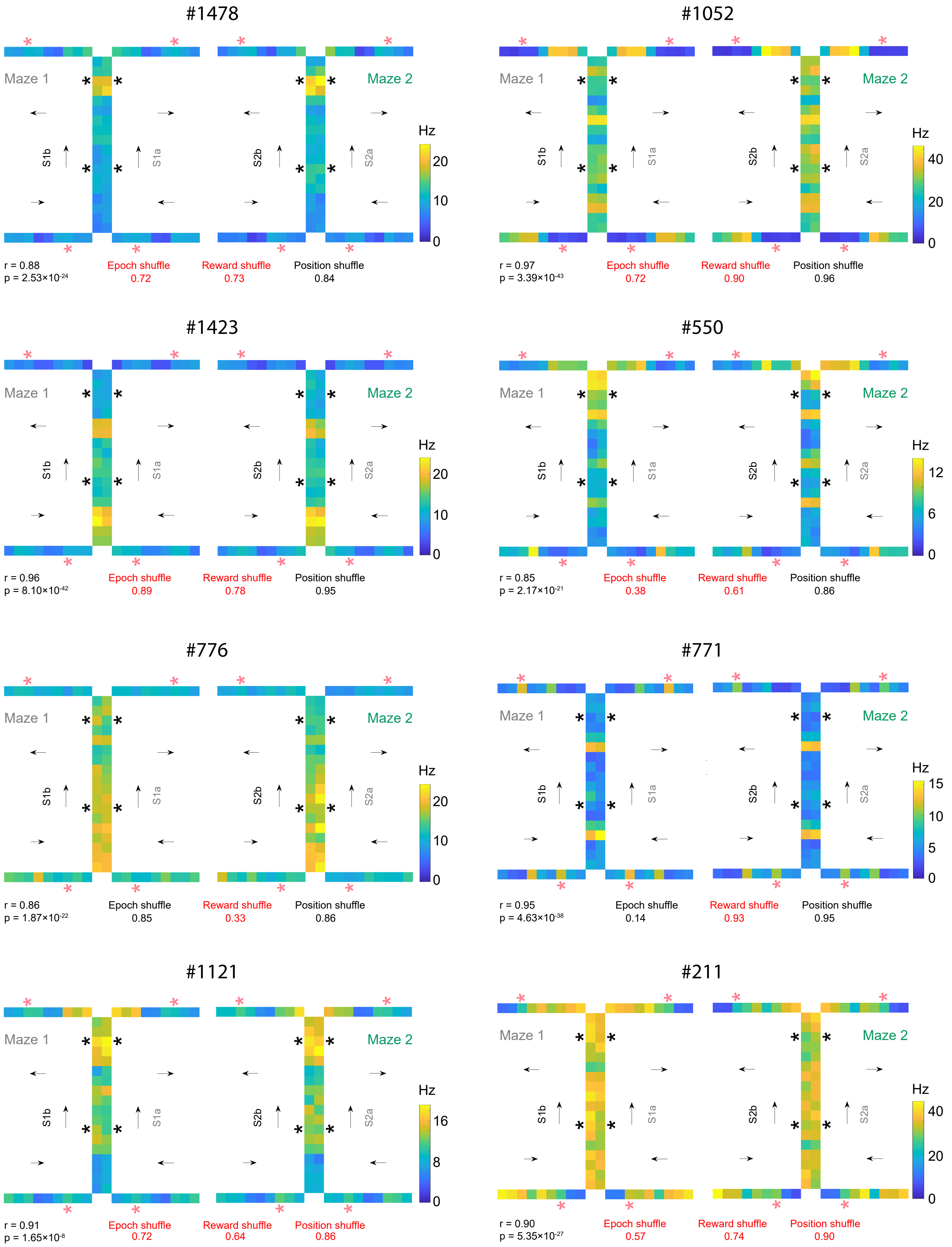
#1478
Maze 1
Maze 2
S1b
S1a
S2b
S2a
Hz
0
10
20
r = 0.88
p = $2.53\times10^{-24}$
Epoch shuffle
0.72
Reward shuffle
0.73
Position shuffle
0.84
#1052
Hz
0
20
40
r = 0.97
p = $3.39\times10^{-43}$
Epoch shuffle
0.72
Reward shuffle
0.90
Position shuffle
0.96
#1423
Hz
0
10
20
r = 0.96
p = $8.10\times10^{-42}$
Epoch shuffle
0.89
Reward shuffle
0.78
Position shuffle
0.95
#550
Hz
0
6
12
r = 0.85
p = $2.17\times10^{-21}$
Epoch shuffle
0.38
Reward shuffle
0.61
Position shuffle
0.86
#776
Hz
0
10
20
r = 0.86
p = $1.87\times10^{-22}$
Epoch shuffle
0.85
Reward shuffle
0.33
Position shuffle
0.86
#771
Hz
0
5
10
15
r = 0.95
p = $4.63\times10^{-38}$
Epoch shuffle
0.14
Reward shuffle
0.93
Position shuffle
0.95
#1121
Hz
0
8
16
r = 0.91
p = $1.65\times10^{-8}$
Epoch shuffle
0.72
Reward shuffle
0.64
Position shuffle
0.86
#211
Hz
0
20
40
r = 0.90
p = $5.35\times10^{-27}$
Epoch shuffle
0.57
Reward shuffle
0.74
Position shuffle
0.90

**Figure S3. Exemplar units illustrate the generalization of positional information across mazes in OFC.**

A pair of heatmaps are shown for each neuron, plotting average activity in each epoch at each position in each maze. Arrows represent sequence directions. Red * marks the reward epoch on rewarded trial types (P3 and P4), while black * marks the reward epoch for non-rewarded trial types (P1 and P2). The correlation coefficients before and after shuffling of epoch, reward, and position are presented in the lower panel of each figure; numbers shown in red were significantly affected by shuffling.

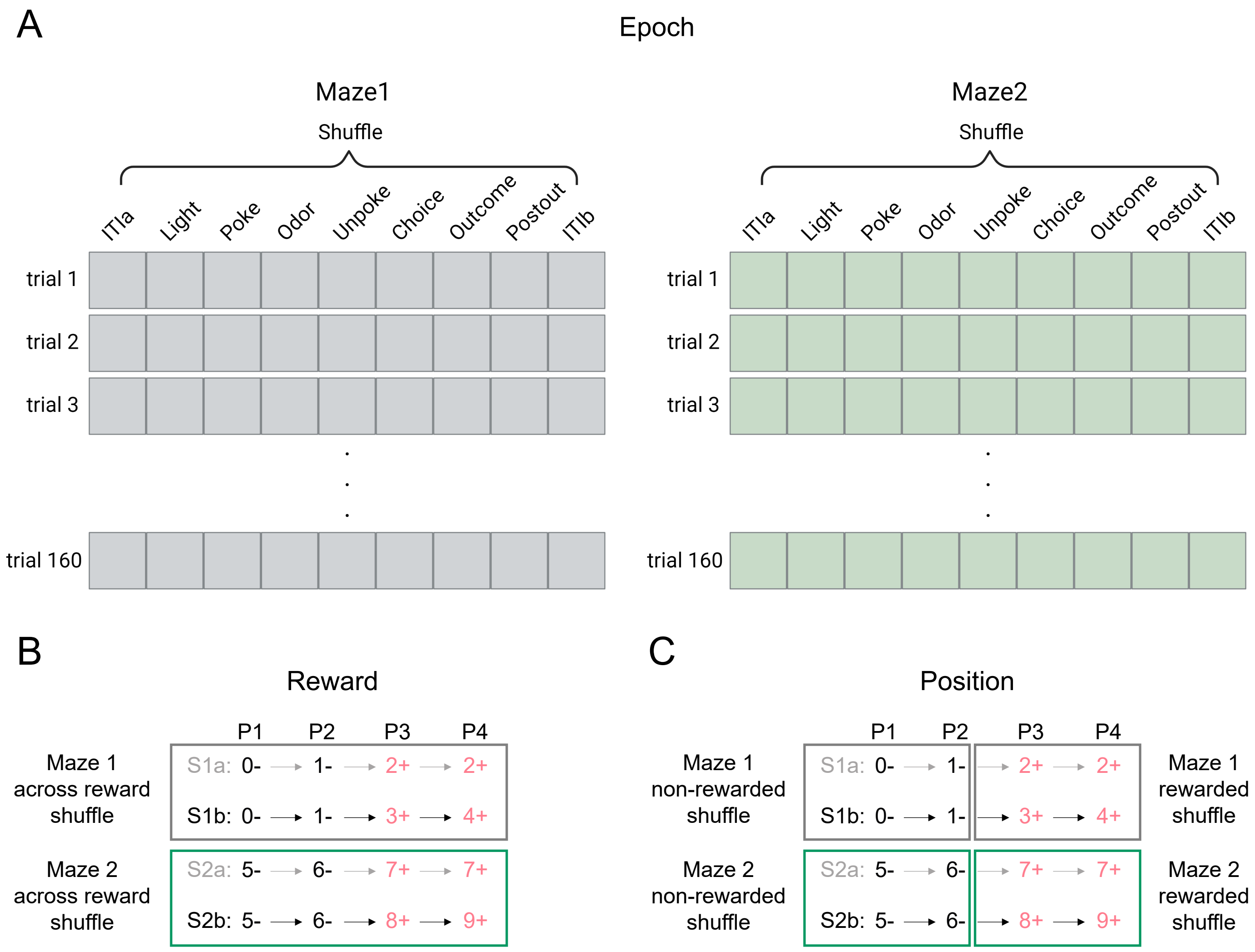


**Figure S4. Illustration of the shuffling used to identify the influence of information related to poch, reward, and position.**

(A) An illustration demonstrating the shuffling of data across all epochs within each maze while maintaining the same position/reward category 1000 times.

(B) Depicts position shuffling by shuffling of data across rewarded and non-rewarded trials within each maze while keeping the epoch constant 1000 times.

(C) Illustration of within-reward position shuffling by shuffling data between positions in each maze but keeping reward category and epoch the same 1000 times. Gray and green colors are used to denote Maze1 and Maze2, respectively.

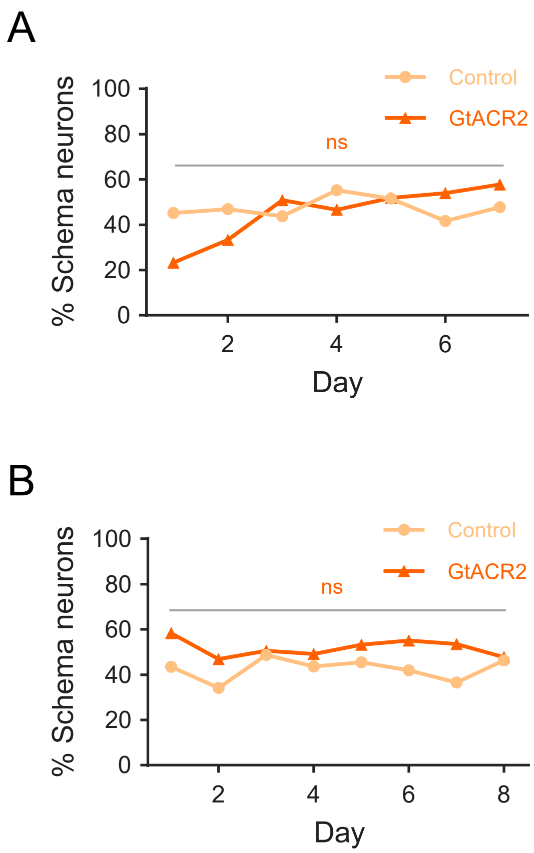


**Figure S5. Percentage of schema neurons during prior training from the rats comprising the control and GtACR2 groups in the learning experiment (Figure 6).**

(A-B) Percentage of schema neurons on each day of retraining after surgery (A) and from control sessions on an established problem (B). There were no differences between the rats in the two groups on any prior day of training (retraining: $\chi^2$'s < 6.0; p's > 0.10; Chi-squared test; well-learned problem: $\chi^2$'s < 3.94; p's > 0.38; Chi-squared test). False discovery rate (FDR) and Benjamini-Hochberg (BH) corrections were applied to correct for multiple comparisons.

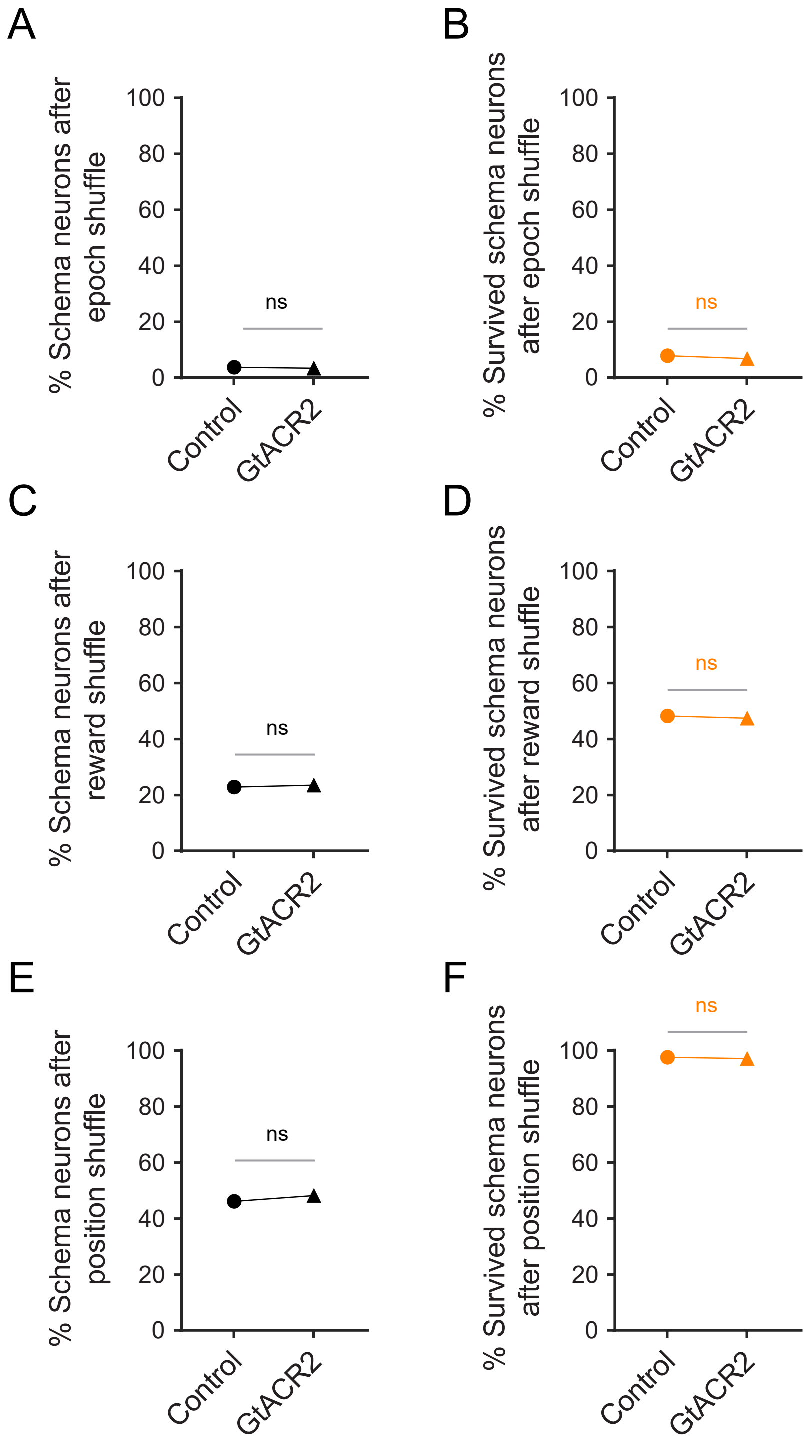


**Figure S6. Effect of ventral subiculum inactivation on percentage of schema cells remaining after shuffling of epoch, reward, and position on an established problem.**

Ventral subiculum inactivation had no effect on the number of remaining schema neurons after shuffling of epoch, reward, or position, whether calculated as a percentage of all neurons (a, c, and e; $\chi^2$'s < 1.68; p's > 0.19) or as a percentage of schema neurons (b, d, and f; $\chi^2$'s < 0.59; p's > 0.44). ns, not significant.

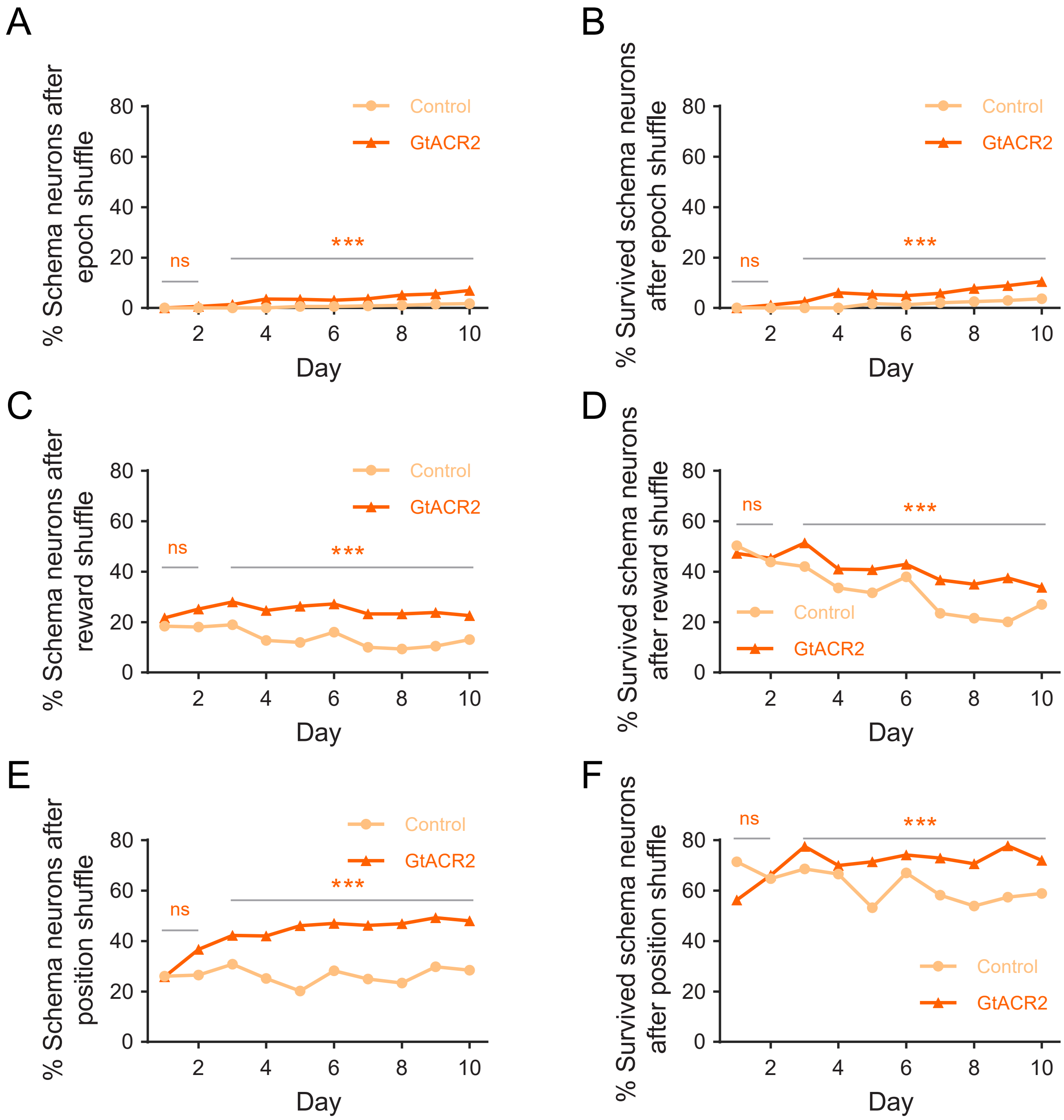

A
% Schema neurons after epoch shuffle
Control
GtACR2
ns
***
Day
B
% Survived schema neurons after epoch shuffle
Control
GtACR2
ns
***
Day
C
% Schema neurons after reward shuffle
Control
GtACR2
ns
***
Day
D
% Survived schema neurons after reward shuffle
Control
GtACR2
ns
***
Day
E
% Schema neurons after position shuffle
Control
GtACR2
ns
***
Day
F
% Survived schema neurons after position shuffle
Control
GtACR2
ns
***
Day

**Figure S7. Effect of ventral subiculum inactivation on percentage of schema cells remaining after shuffling of epoch, reward, and position during learning of a new problem.**

Ventral subiculum inactivation accelerated the formation of schema neurons (Figure 5C), resulting in a significantly larger number of remaining schema cells in the inactivated rats after shuffling of epoch, reward or position, whether represented as the proportion of all neurons (A, C, and E) or as the percentage of schema neurons (B, D, and F).

(A-B) Chi-squared tests: (A, B) Quantification of the percentage of schema neurons after epoch shuffle on each day against all neurons(A) (Overall: $\chi^2 = 72.9$, $p = 1.4 \times 10^{-17}$; days 1-2: $\chi^2 = 1.4$; $p = 0.23$; days 3-10: $\chi^2 = 77.2$, $p = 1.6 \times 10^{-18}$; Chi-squared test) and against schema neurons (B) before epoch shuffle (Overall: $\chi^2 = 10.0$, $p = 0.0016$; days 1-2: $\chi^2 = 1.2$; $p = 0.26$; days 3-10: $\chi^2 = 16.4$, $p = 5.2 \times 10^{-5}$; Chi-squared test). The groups were similar in the beginning but showed divergence over time.

(C-D) The percentage of schema neurons compared to all neurons(C) (Overall: $\chi 2 = 19.7$, $p = 9.0 \times 10^{-6}$; days 1-2: $\chi 2 = 0.58$, $p = 0.45$; days 3-10: $\chi 2 = 18.1$, $p = 2.1 \times 10^{-5}$; Chi-squared test) and original schema neurons(D) (Overall: $\chi 2 = 11.5$, $p = 6.9 \times 10^{-4}$; days 1-2: $\chi 2 = 0.44$, $p = 0.51$; days 3-10: $\chi 2 = 10.2$, $p = 0.0014$; Chi-squared test) were quantified on a daily basis after reward shuffle.

(E-F) The percentage of schema neurons relative to all neurons(E) (Overall: $\chi^2 = 43.7$, $p = 3.9 \times 10^{-11}$; days 1-2: $\chi^2 = 2.4$; $p = 0.12$; days 3-10: $\chi^2 = 44.8$, $p = 7.7 \times 10^{-4}$; Chi-squared test) and total schema neurons before position shuffle (F) (Overall: $\chi^2 = 7.8$, $p = 0.005$; days 1-2: $\chi^2 = 0.0032$; $p = 0.96$; days 3-10: $\chi^2 = 11.3$, $p = 7.7 \times 10^{-4}$; Chi-squared test) were measured for each day following position shuffle. ***$p < 0.001$; ns, not significant.

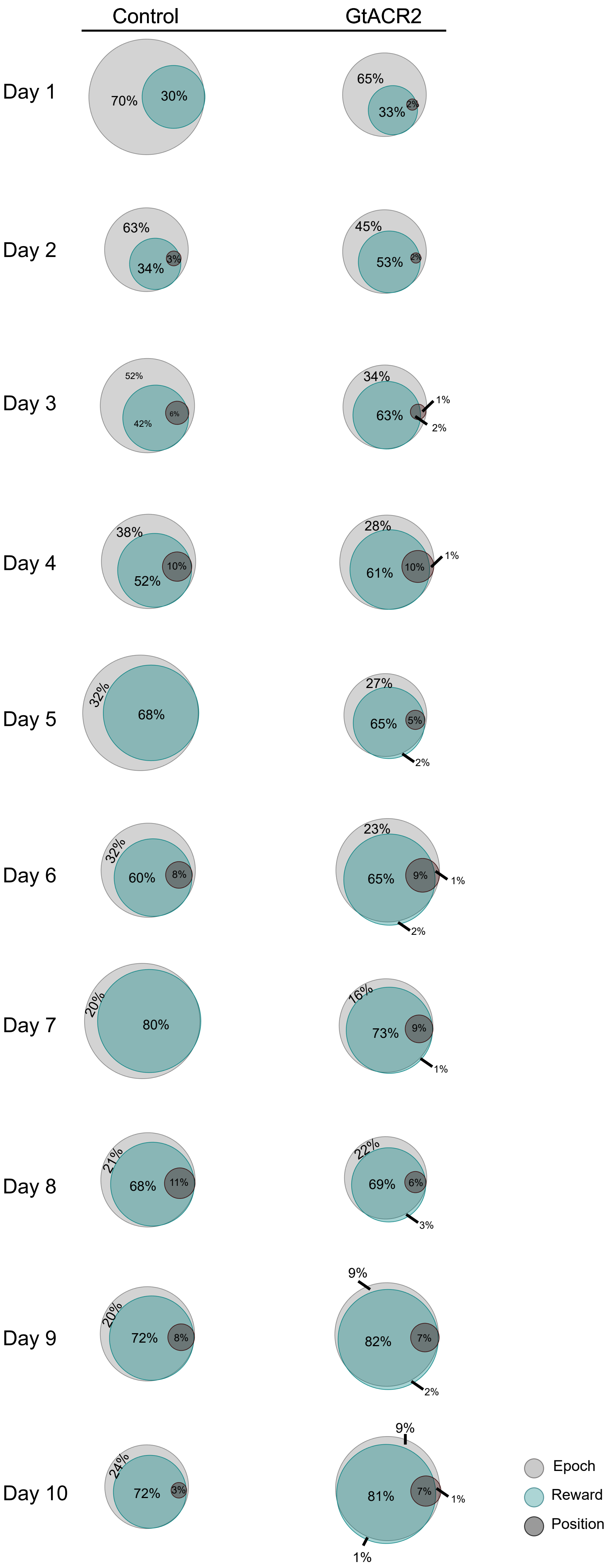
Control
GtACR2
Day 1
70%
30%
65%
33%
2%
Day 2
63%
34%
3%
45%
53%
2%
Day 3
52%
42%
6%
34%
63%
1%
2%
Day 4
38%
52%
10%
28%
61%
10%
1%
Day 5
32%
68%
27%
65%
5%
2%
Day 6
32%
60%
8%
23%
65%
9%
1%
2%
Day 7
20%
80%
16%
73%
9%
1%
Day 8
21%
68%
11%
22%
69%
6%
3%
Day 9
20%
72%
8%
9%
82%
7%
2%
Day 10
24%
72%
3%
9%
81%
7%
1%
1%
Epoch
Reward
Position

**Figure S8. Effect of ventral subiculum inactivation on the proportions of schema neurons impacted by shuffling of epoch, reward, and position-related information by day.**

Venn diagrams summarize data from Figure 7A-D, which display the percentage of schema neurons that were recorded during control and GtACR2 sessions and how they were influenced by information related to epoch (light gray), reward (light green), and position (dark gray) across days.
